# A generative framework for enhanced cell-type specificity in rationally designed mRNAs

**DOI:** 10.1101/2024.12.31.630783

**Authors:** Matvei Khoroshkin, Arsenii Zinkevich, Elizaveta Aristova, Hassan Yousefi, Sean B. Lee, Tabea Mittmann, Karoline Manegold, Dmitry Penzar, David R. Raleigh, Ivan V. Kulakovskiy, Hani Goodarzi

## Abstract

mRNA delivery offers new opportunities for disease treatment by directing cells to produce therapeutic proteins. However, designing highly stable mRNAs with programmable cell type-specificity remains a challenge. To address this, we measured the regulatory activity of 60,000 5’ and 3’ untranslated regions (UTRs) across six cell types and developed PARADE (Prediction And RAtional DEsign of mRNA UTRs), a generative AI framework to engineer untranslated RNA regions with tailored cell type-specific activity. We validated PARADE by testing 15,800 de novo-designed sequences across these cell lines and identified many sequences that demonstrated superior specificity and activity compared to existing RNA therapeutics. mRNAs with PARADE-engineered UTRs also exhibited robust tissue-specific activity in animal models, achieving selective expression in the liver and spleen. We also leveraged PARADE to enhance mRNA stability, significantly increasing protein output and therapeutic durability in vivo. These advancements translated to notable increases in therapeutic efficacy, as PARADE-designed UTRs in oncosuppressor mRNAs, namely PTEN and P16, effectively reduced tumor growth in patient-derived neuroglioma xenograft models and orthotopic mouse models. Collectively, these findings establish PARADE as a versatile platform for designing safer, more precise, and highly stable mRNA therapies.

## Introduction

mRNA therapeutics are revolutionizing disease treatment by leveraging messenger RNA (mRNA) to direct cells to produce therapeutic proteins, offering capabilities beyond those of traditional small molecules^1^. However, despite significant advances, current mRNA therapeutics face critical challenges, particularly in achieving stability and cell type-specificity of therapeutic mRNAs. Rapid degradation of mRNA molecules compromises their efficacy^2^, while insufficient cell type-specificity may produce off-target effects, causing toxicity in non-relevant tissues^3^.

Cell type-specificity at the transcriptional level has been extensively studied and successfully achieved, aiding the development of DNA-based therapies^4,5^. Yet, the post-transcriptional regulation, crucial for mRNA therapeutics, remains less explored^6–8^, and designing mRNAs with cell type-specific expression, i.e., preferential activity in certain cell types over others, remains challenging^9,10^. One common strategy involves incorporating microRNA binding sites into the 3’ untranslated region (3’UTR), utilizing cell type-specific microRNAs to selectively suppress mRNA activity^11^. However, this approach has shown limited success, as it fails to generalize across cell types due to the variability in microRNA expression^12^ and activity.

A critical hurdle in mRNA design lies in the lack of large-scale and consistently measured datasets needed to study post-transcriptional regulation in different cell types. Most machine learning models aimed at optimizing mRNA activity, such as enhancing translation efficiency or stability, either rely solely on RNA sequence data or are trained on measurements from a single cell line, overlooking the heterogeneity of cell types^13–16^. The few models that do account for cell type-specificity suffer from dataset heterogeneity and biases introduced by varying experimental protocols. For example, ribosome profiling (Ribo-Seq) is suitable for assessing ribosome occupancy, yet these measurements are limited to a few thousand genes and are often influenced not only by the mRNA sequence but also by transcriptional and co-transcriptional effects at individual genomic loci^17^, introducing undesired variance and complicating the identification of post-transcriptional regulatory signals within the sequence. Furthermore, large-scale collections of Ribo-Seq data span datasets collected across different laboratories using various non-standardized protocols, further contributing to heterogeneity^18^. Although deep learning has shown promise in related areas, such as designing transcriptional enhancers^19–21^, the absence of suitable data has limited its application to cell type-specific mRNA design^22^. Similarly, while a plethora of datasets measuring static RNA levels using RNA-Seq is available, these datasets are confounded by transcriptional activity^23^, making it difficult to discern the sequence determinants of post-transcriptional regulation.

To address these issues, we used massively parallel reporter assays (MPRA)^24^ to directly measure the influence of untranslated regions (both 5’UTRs and 3’UTRs) on RNA activity—specifically, the amount of protein produced from a given RNA—avoiding the confounders introduced at the transcriptional level. These MPRAs yielded a large-scale dataset covering multiple cell types, and provided a clearer view of post-transcriptional regulation by focusing solely on the impact of untranslated regions. This dataset enabled us to develop PARADE (Prediction And RAtional DEsign of RNA UTRs), a comprehensive artificial intelligence (AI) framework that combines predictive and generative capabilities to optimize UTR sequences for cell type-specific mRNA stability and translation. PARADE includes two primary modules: the PARADE Predictor, which accurately forecasts UTR activity and specificity across cell types, and the PARADE Generator, which designs synthetic UTR sequences with tailored activity profiles. Together, these modules enable PARADE to generate synthetic RNA sequences that demonstrate superior cell type-specificity and stability compared to naturally occurring sequences and existing RNA therapeutics.

We further demonstrated the practical application of the PARADE Framework by designing mRNAs that are selectively active in different cell types. In particular, we engineered UTRs for mRNA encoding the hepatotoxic gene product CYP2E1 to be active in T cells but not in hepatocytes, effectively reducing hepatotoxicity. In animal models, mRNAs designed by the PARADE Generator displayed distinct activity profiles, achieving selective expression in hepatocyte-enriched and T cell-enriched tissues such as liver and spleen. Additionally, we used the PARADE Generator to design UTR sequences with prolonged activity that enabled the suppression of neuroglioma xenografts and orthotopic models in mice by delivering PTEN- or P16-encoding mRNA. PARADE Framework is generalizable and can be applied to design mRNA UTRs selectively active in a variety of cell types, providing a powerful tool for developing safer, more effective RNA-based therapeutics.

## Results

### Massively Parallel Reporter Assays Provide a Large-Scale Dataset of Cell Type-Specific UTR Activity

To assess the effect of UTRs on post-transcriptional regulation, in isolation and at scale, we selected 60,000 UTR fragments from the human transcriptome and evaluated their impact on reporter gene expression in six different cell lines. We began by identifying 2068 transcripts with cell type-related variability in translation efficiency from published Ribo-Seq data^25–27^ as we hypothesized that these transcripts likely harbor regulatory elements driving differential activity patterns across cell types.

From this set of transcripts, we randomly sampled 60,000 regions from their respective 5’ and 3’ UTRs, creating a diverse sequence library, which we refer to as Natural Set (Library 1). Next, we performed MPRAs to evaluate the activity of these UTR segments across six cell lines, estimating protein output from genome-integrated reporter constructs via flow cytometry^28^.

For the 5’ UTRs, we constructed a library of 30,000 segments (50 nt long), which we cloned upstream of the eGFP open reading frame (ORF) in a polycistronic eGFP-mCherry reporter (Fig. 1A). In this design, eGFP and mCherry are both transcribed from a single promoter but are translated separately: eGFP in a cap-dependent manner and mCherry in a cap-independent manner.

**Figure 1.**
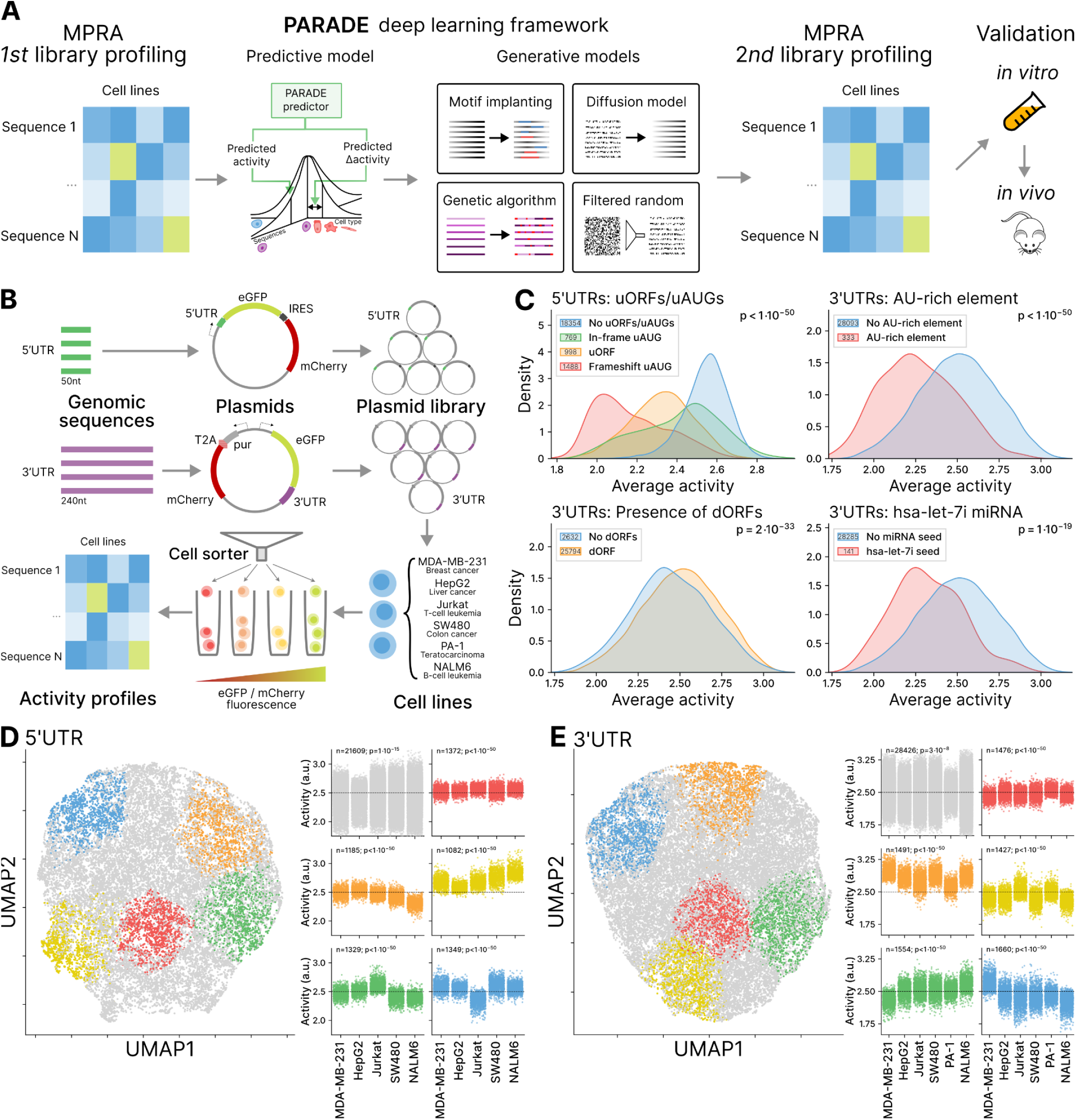
Massively Parallel Reporter Assay Reveals Cell Type-Specific UTR Activity Patterns. **A.** Overview of the PARADE Framework workflow from MPRA profiling to deep learning model, generative sequence design, and experimental validation. **B.** Schematic of the MPRA experiment. The reporter construct includes an internal ribosome entry site (IRES) for cap-independent translation, a T2A sequence enabling co-expression of the reporter and selection marker, and a puromycin resistance gene (pur) for selecting successfully transfected cells. **C.** Effects of known regulatory elements on UTR activity. The kernel density estimate plot shows the relationship between the average activity and the presence of upstream open reading frames (uORFs/uAUGs; top left), downstream open reading frames (dORFs; bottom left), AU-rich elements (top right), and selected miRNA seed sequences (bottom right). P-values: Kruskal - Wallis nonparametric test. **D, E.** UMAP visualization of cell type-specific activity profiles for 5’UTRs (D) and 3’UTRs (E). Each point (left) represents a UTR sequence, colored by K-means clustering of activity profiles across the cell types. The activity levels for colored clusters are shown on the right, grey points denote the rest of the UTRs. p-values: Kruskal - Wallis nonparametric test.

Importantly, this design allows us to decouple the effects of 5’ UTR sequences on translation from those on transcription^29^. While variations in the 5’ UTR sequences can impact the translation of eGFP, they do not affect the cap-independent translation of mCherry, providing an internal control. We transduced these libraries into six cell lines representing different tissues: Jurkat (T cells), Nalm-6 (B cells), SW-480 (colon), PA-1 (ovary), MDA-MB-231 (breast), and HepG2 (liver). We then sorted the cells from each library into four bins by eGFP/mCherry expression ratios and sequenced the DNA from the sorted pools to identify respective UTRs and analyze their activity. Throughout this study, we define sequence “activityˮ as the relative expression of the reporter gene, measured by the “center of massˮ of normalized read counts across the four cell sorting bins.

For the 3’ UTRs, we constructed a sister library of 30,000 segments (240 nt long), which were cloned downstream of the eGFP ORF in a bidirectional eGFP-mCherry reporter (Fig. 1B). In this setup, eGFP and mCherry were produced from separate RNAs, thus the changes in the eGFP/mCherry ratio reflected how the 3’ UTRs influenced both translation and RNA stability. As with the 5’ UTRs, we sorted the cells into four groups based on their eGFP/mCherry expression ratio, followed by sequencing, read counting, and normalization.

To confirm the reliability of our measurements, we evaluated whether known regulatory elements in the 5’ and 3’ UTRs acted as expected (Fig. 1C). First, in the 5’ UTRs, we analyzed the effects of upstream open reading frames (uORFs) known to repress translation by interfering with ribosome scanning and preventing translation initiation on the primary ORF^30^. Indeed, the presence of uORFs decreased translation activity; notably, the presence of frame-shifted upstream start codons (uAUGs) had a stronger impact on activity compared to the in-frame uAUG (p<10^-50^).

For 3’ UTRs, downstream open reading frames (dORFs) in the 3’UTRs slightly but significantly (p=2:10^-33^) enhanced the activity of the main ORF as shown previously^31^. The presence of AU-rich elements (AREs), associated with mRNA instability and decreased translation^32^, on the other hand, decreased the sequence activity in our MPRAs (p<10^-50^). Lastly, we assessed the influence of the hsa-let-7i miRNA, a known post-transcriptional repressor^33^, and observed a corresponding reduction in average activity for sequences containing its seed sequence (p=2:10^-19^). These observations confirm the validity of our approach not only in capturing the regulatory effects of specific elements but also in quantitatively and broadly measuring the effects of RNA sequence on the post-transcriptional regulation.

We next examined whether the MPRA data revealed UTRs with cell type-specific activity patterns across the tested cell lines. While previous studies have shown that 3’UTRs harbor cell type-specific regulatory elements^34^, 5’ UTRs have been considered less likely to contribute to cell type-specific expression patterns^35^. In our data, the activity levels of individual sequences were generally well-correlated across the cell lines. Notably, Pearson correlation observed between cell lines was, on average, 25% lower than between replicates, with median *r* values of 0.636 for 5’ UTRs and 0.619 for 3’ UTRs for cell lines, compared to 0.856 and 0.820 for replicates (Suppl. Fig. 1B). Despite the overall correlation, we sought to identify the sequences that demonstrated differential activity across the cell lines. For this, we calculated the *τ* index^36^ which is a robust metric for tissue specificity; *τ* ranges from 0 to 1, where 0 represents ubiquitous expression (housekeeping genes) and 1 indicates exclusive expression in a single cell type. Using 0.2 is a cut-off, we identified 4897 3’ UTRs and 412 5’ UTRs (out of ∼30,000 in each respective set) with varied patterns of activity across the cell lines.

To further illustrate these patterns of cell type-specificity, we applied unbiased K-means clustering to the sequence activity measurements across the tested cell lines. This analysis revealed distinct clusters of sequences with non-uniform activity profiles (Fig. 1D and 1E). Typical cell type-specific activity patterns included sequences with higher or lower activity in Jurkat and PA-1 compared to other cell lines for 3’ UTRs, represented by the yellow and orange clusters, respectively. For 5’ UTRs, a distinct yellow cluster captured sequences with higher activity in Nalm-6 and SW480 but lower activity in other cell lines. Additional examples included sequences with higher activity in MDA-MB-231 for 3’ UTRs, forming the blue cluster, and sequences with higher or lower activity in Jurkat for 5’ UTRs, represented by the green and blue clusters, respectively. Importantly, cell type-specific activity was observed for both 3’ and 5’ UTRs, suggesting that determinants of cell type-specific post-transcriptional regulation are present in both regions. Collectively, our high-throughput MPRA data are consistent with the presence of known regulatory elements in UTRs and capture the cell type-specific activity profiles of a large set of sequences.

### PARADE accurately predicts the activity of 5’ and 3’ UTRs and accounts for key regulatory elements

To reveal the sequence determinants of cell type-specificity, we first devised and trained a deep learning model, PARADE Predictor, using our large MPRA dataset. PARADE Predictor is based on the LegNet architecture, which has previously demonstrated excellent performance in predicting DNA regulatory activity from transcriptional MPRA data^37^. PARADE Predictor accurately estimated both the absolute activity of a given sequence in a specific cell line as well as its “activity deviation” Δ, i.e. the difference between its activity in the query cell line and the average activity across all lines (Fig. 2A, Extended Data Fig. 2A). In these tasks, PARADE Predictor outperformed both the regression with k-mer counts and Optimus-5-prime^38^, the state-of-the-art model for 5’ UTRs, across all cell lines (Fig. 2B), estimating cell type-specific sequence activity with Pearson correlation *r* of 0.65-0.79 for 5’ UTRs and 0.65-0.75 for 3’ UTRs. The model performance was improved by including additional features (Extended Data Fig. 2D-E). As expected, inclusion of the triplet phase annotation improved PARADE Predictor’s performance for 5’ UTRs, but did not affect 3’ UTRs. Training PARADE Predictor to estimate both the average activity and deviations from this average in each line (i.e. activity deviation Δ) resulted in higher correlation scores for 3’ but not 5’ UTRs.

**Figure 2.**
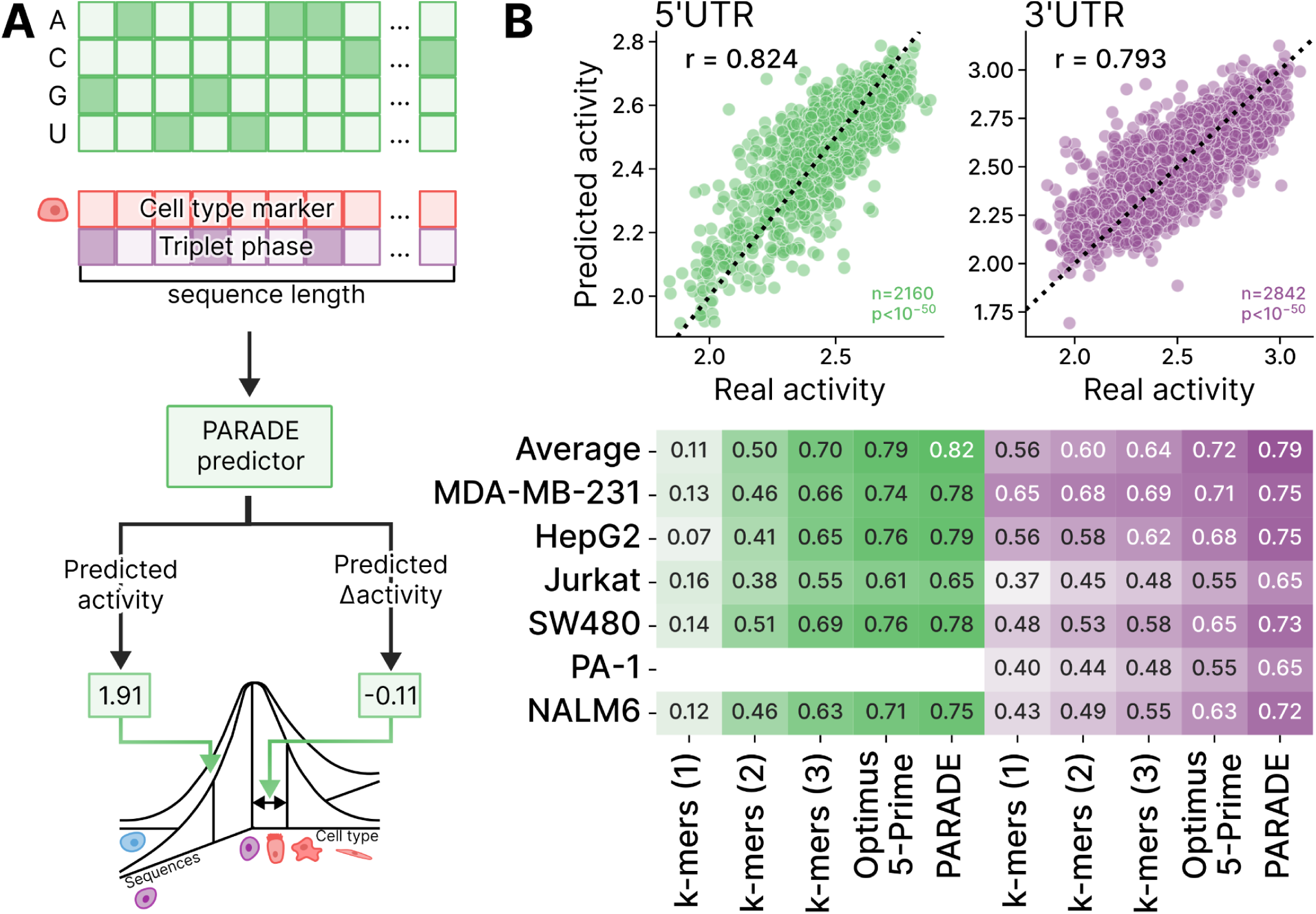
PARADE Predictor Accurately Estimates UTR Activity and Cell Type-Specificity. **A.** Schematic of the PARADE Predictor: a LegNet-based neural network is trained on one-hot encoded UTR sequences with triplet phase and cell type provided in extra channels. The model outputs both the absolute sequence activity in specific cell lines and the activity deviation (Δ), the difference from the average activity across all cell lines. **B.** Top: Scatterplots comparing PARADE Predictor’s estimates of average activity for 5’UTRs (green) and 3’UTRs (purple) with the experimentally measured values. Bottom: Pearson correlation (r) of predicted and experimentally measured activity for PARADE Predictor and other models across different cell lines; the color scale saturation also denotes r.

To evaluate PARADE Predictor’s ability to estimate the cell type-specific activity profiles, we focused on two metrics: the cell type-specificity index *τ*^36^ and the “activity deviation” Δ (see above). Considering cell type-specific predictions, PARADE Predictor more accurately approximated the true *τ* index for individual sequences, showing a 30% and 50% improvement over Optimus-5-prime for 5’ UTRs and 3’ UTRs, respectively (Pearson correlation *r* of 0.31 vs. 0.24 for 5’ UTRs and 0.39 vs. 0.26 for 3’ UTRs, Extended Data Fig. 2B). Additionally, PARADE Predictor achieved higher accuracy in evaluating Δ than both Optimus-5-prime and k-mer regression in every single cell line (Extended Data Fig. 2C). Overall, these results demonstrate PARADE Predictor’s superior accuracy in forecasting both the average and the cell type-specific activity.

### PARADE Identifies The Key Motifs Contributing to Cell Type-Specific Activity

To uncover the sequence grammar driving the cell type-specific UTR activity, we performed motif analysis using two complementary approaches. First, we conducted motif discovery directly from the MPRA data using FIRE^39^. Second, we analyzed the sequence patterns learned by PARADE Predictor using TF–MoDISco^40^. While TF–MoDISco did not find any significant motifs in the 5’ UTRs, both methods converged on a similar set of motifs in the 3’ UTRs. We annotated the discovered motifs by comparing them to those of known RNA-binding proteins (RBPs) (Fig. 3A, Extended Data Fig. 3) and revealed both well-characterized regulators of translation control and mRNA stability and potentially novel regulatory elements, such as CUGCMW in 3’ UTRs (Fig. 3B).

**Figure 3.**
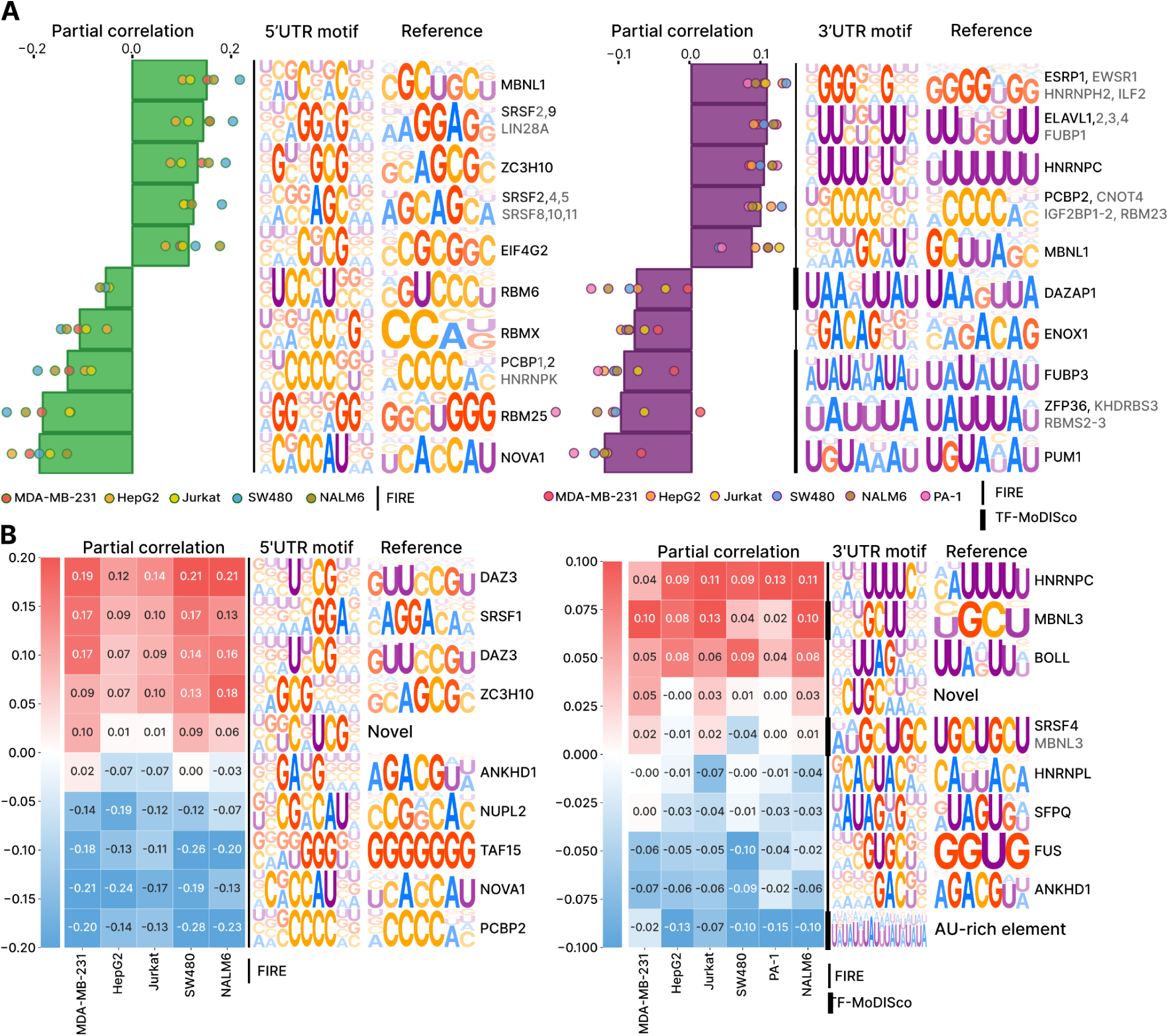
PARADE Predictor Identifies Key Motifs of RNA-Binding Proteins Governing the Cell Type-Specificity. **A.** Ten motifs with the highest absolute mean partial correlation between motif scores and experimentally measured sequence activity. RNA-binding protein motifs were identified using FIRE and TF-MoDISco, and clustered by similarity to known motifs from the oRNAment and CIS-BP-RNA databases to derive representative reference motifs. Bars represent the mean partial correlation (corrected for the nucleotide composition), and individual colored dots correspond to partial correlations in specific cell lines (green for 5’UTRs, purple for 3’UTRs). **B.** Heatmap of partial correlations for selected motifs that demonstrate variable activity across different cell lines, highlighting cell type-specific regulatory effects.

To assess the varying influence of RBP binding motifs across cell lines, we calculated partial correlations between motif scores and experimentally measured sequence activities, explicitly accounting for nucleotide composition as a confounding factor (see Methods). We found that for many, though not all, RBP binding motifs, their influence on UTR activity varied across cell lines (Fig. 3A,B). For instance, in the 3’ UTR, DAZAP1 is linked to strong negative regulation in most cell lines, but shows minimal impact in breast cancer MDA-MB-231 cells. This observation aligns with previous studies, which showed that DAZAP1 depending on the context can participate in both mRNA degradation and silencing^41^ as well as in activation of mRNA translation^42^. Similarly, MBNL1 exhibits positive regulation of 3’UTRs in lymphocytes (Jurkat and Nalm-6) and hepatocytes (HepG2), but its effect is minimal in ovarian PA-1 and colon SW480 cells.

Cell type-specific effects of regulatory motifs are likely mediated by variations in the expression or binding activity of their cognate RNA-binding proteins. For example, presence of motif occurrences of PUM1, a key driver of ovarian cancer^43^, correlates with lower 3’UTR activity across cell lines, with the strongest effect in the ovarian cancer cell line PA-1. Similarly, the motif occurrences of SRSF9, which stabilizes mRNA in colorectal cancer by interacting with m6A-methylated regions^44^, are correlated with 5’ UTR activity, particularly in the colorectal cancer (SW480) cell line. These examples underscore PARADE Predictor’s utility in capturing cell type-specific regulatory mechanisms associated with well-known RBPs and novel regulatory elements.

### PARADE Generator Expands Sequence Diversity and Achieves High Cell Type-Specificity

From the start, we devised PARADE Framework to go beyond prediction and integrate generative capabilities to design UTR sequences with predefined activity levels and specificity. While PARADE Predictor assesses the activity of candidate UTR sequences, PARADE Generator employs several generative methods — Diffusion, Genetic Algorithm, Random Sampling, and Motif-based design — to create sequences that meet these predefined activity levels and specificity requirements (Fig. 4A,B). These methods aim to maximize the UTR’s Cell Type Activity Difference (CTAD) — a measure of specificity defined as the difference in activity between two selected cell lines, where higher absolute values indicate greater specificity.

**Figure 4.**
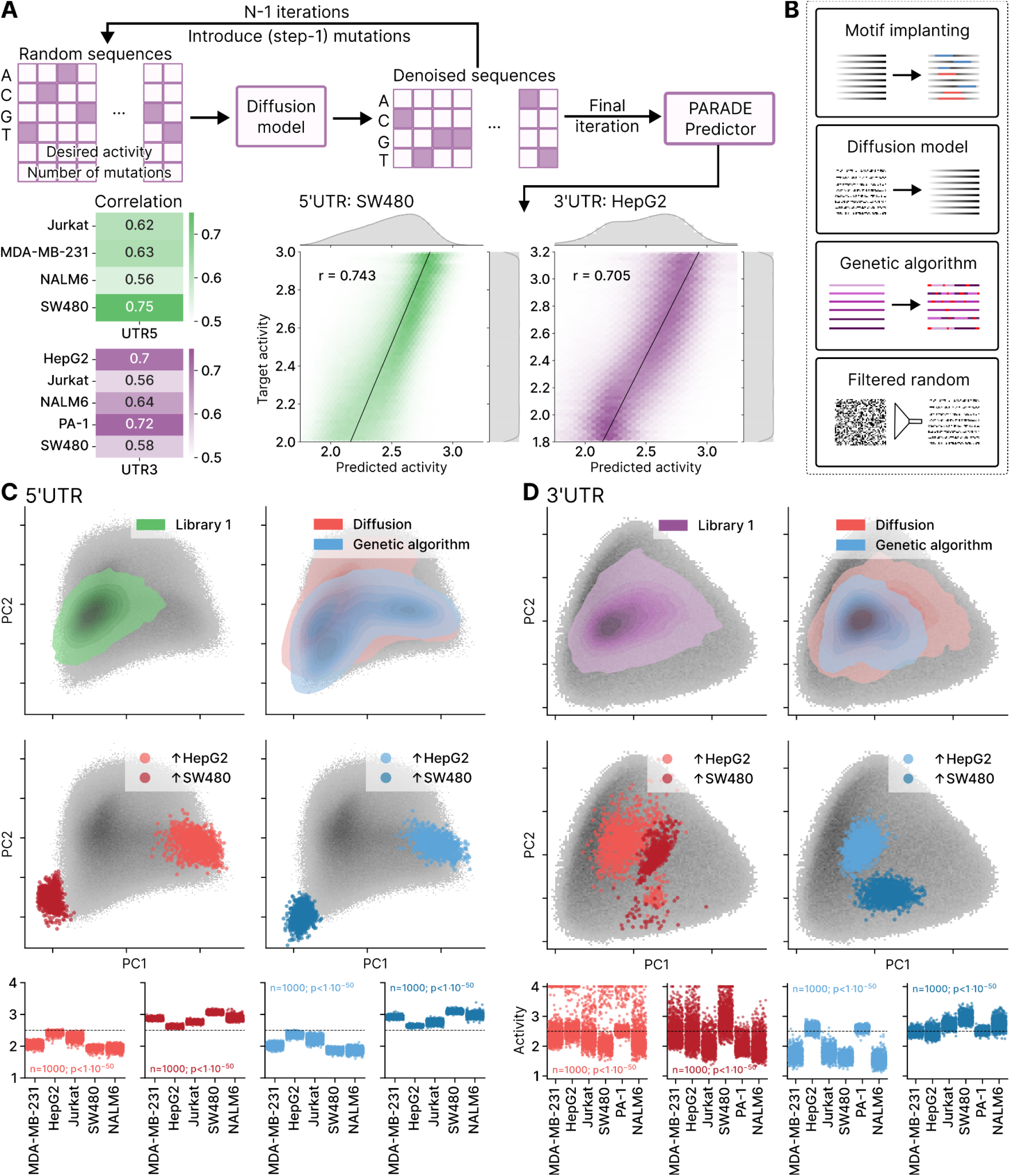
PARADE Generator Produces Diverse and Highly Cell Type-Specific UTR Sequences. **A.** Schematic of the Diffusion model for sequence generation. Top: Workflow overview. Bottom left: Achieved correlations of the target activity and the resulting (predicted) activity across the cell types. Bottom right: Hexagonally-binned two-dimensional histograms of the target versus the predicted activity for 5’UTRs in SW480 and 3’UTRs in HepG2. **B.** Overview of the four sequence generation methods comprising the PARADE Generator. Diffusion iteratively refines random sequences through mutations to achieve desired activity profiles. Genetic Algorithm refines sequences by iteratively applying selection, mutation, and crossover processes to maximize cell-specific regulatory activity. The motif-based design combines known regulatory motifs into novel configurations to drive cell-specific activity. Random sampling generates sequences by shuffling nucleotides and selecting those with the highest predicted CTAD. Detailed plots illustrating these methods are provided in Extended Data Figure 4A-D. **C.** Visualization of the latent space of the 5’UTR PARADE Framework: PCA of the model embeddings for Library 1 (top left) and sequences generated by the diffusion model and genetic algorithm (top right). Random sequences are shown in grey as background. Sequences with the highest Cell Type Activity Difference (CTAD) for HepG2 and SW480 are highlighted for Diffusion (middle left) and Genetic algorithm (middle right). Bottom: Predicted activity for highlighted sequences. p-values: Kruskal-Wallis nonparametric test. **D.** Visualization of the model embeddings from the 3’UTR PARADE Framework with the same approach as in panel C.

The first approach within the PARADE Generator was Diffusion, a generative model based on the cold diffusion principle^45^. Random sequences were iteratively refined through a series of mutations to shift them through the model’s latent space closer to the subspace occupied by sequences with the target activity. These refined sequences were then evaluated by the PARADE Predictor, and those with the highest CTADs were taken for further evaluation. Diffusion achieved a strong correlation between the preset and predicted activity levels, ranging from 0.56 to 0.72 for both 5’ and 3’ UTRs (Fig. 4A). Notably, over 99.8% of the sequences generated by Diffusion were unique and distinct from the training set, demonstrating the model’s capability to design novel sequences^46^.

Next, we implemented a genetic algorithm^47^ that directly optimized CTAD by refining sequences to maximize the differential activity between two cell lines (which we refer to as Genetic Algorithm; Extended Data Fig. 4C). We also employed a random sampling approach (referred to as Random), generating 2 million sequences for both 5’- and 3’UTRs by randomly shuffling sequences from Library 1 while preserving either mono- or di-nucleotide content. The PARADE Predictor was used to identify and select the sequences with the highest CTAD (Extended Data Fig. 4B). Additionally, we used a motif-based design strategy (referred to as Motifs), generating sequences by combining occurrences of two previously identified motifs for 5’ UTRs and four for 3’ UTRs (Extended Data Fig. 4A).

To compare and benchmark these methods, we included two control groups of UTRs. The first control group consisted of characterized sequences, including those from established RNA therapeutics, such as the UTRs used in Moderna’s mRNA-1273 and Pfizer’s BNT162b2 COVID vaccines. For clarity, these vaccine-derived UTRs will be referred to throughout this study as Reference UTR 1 and Reference UTR 2, respectively. This control group also contained a variety of well-characterized regulatory elements to provide a comprehensive baseline for evaluating cell type-specificity^10,48–50^. The second control group contained sequences with combinations of either activating or repressing motifs discovered in our motif analysis.

We hypothesized and subsequently demonstrated that PARADE’s generative AI achieves higher cell type-specificity by designing sequences unconstrained by evolutionary constraints and selection pressure. To test this hypothesis, we applied linear dimensionality reduction (PCA) to the PARADE embeddings of 1 million random sequences (Extended Data Fig. 4D). In this space, we overlaid naturally occurring UTRs from Library 1 alongside sequences generated by Diffusion and Genetic Algorithm. For short 5’ UTRs (50 nts), each of the designed sequence sets occupied a larger area of the latent space compared to the natural sequences, indicating the models’ power in exploring the available sequence space (Fig. 4C). For longer 3’ UTRs (240 nts), the area explored by Genetic Algorithm did not exceed the ’native’ space (Fig. 4D), but Diffusion-generated sequences were again more diverse as for 5’ UTRs.

We also hypothesized that groups of sequences optimized for high CTAD between specific pairs of cell lines would cluster into distinct regions of the latent space. Indeed, these CTAD-optimized sequences occupied distinct regions of the space (Fig. 4C and Fig. 4D). Notably, in some instances, different generative algorithms explored divergent regions within this space (Fig. 4D), suggesting there are alternative regulatory elements and pathways to achieve the desired cell type-specificity.

### Designed RNA Sequences Demonstrate Enhanced Cell Type-Specificity

From sequence sets constructed with 4 generative methods — Diffusion, Genetic Algorithm, Random, and Motifs — and two control groups (characterized sequences and combinations of activating or repressing motif occurrences) — we selected 12,000 sequences for 5’ UTRs and another 12,000 sequences for 3’ UTRs (see Methods). This set of sequences, referred to as Designed Set (Library 2), was then tested using the MPRA assay in the same six cell lines (Jurkat, Nalm-6, SW-480, PA-1, MDA-MB-231, and HepG2) as used for Library 1. Of these, 10,174 and 5,675 sequences from the 5’ and 3’ UTR libraries, respectively, passed the experimental quality control thresholds. Compared to Library 1, Library 2 covered a larger area of the PARADE embedding space (Fig. 5A), indicating a greater diversity of sequences. The purpose of the evaluation was to assess four aspects: (1) the accuracy of PARADE Predictor in evaluating sequence activity, (2) how closely the Diffusion method achieves the desired activity, (3) the level of improvement in the cell type-specificity compared to the controls and Library 1, and (4) the comparative performance of the generative methods in enhancing the cell type-specificity.

**Figure 5.**
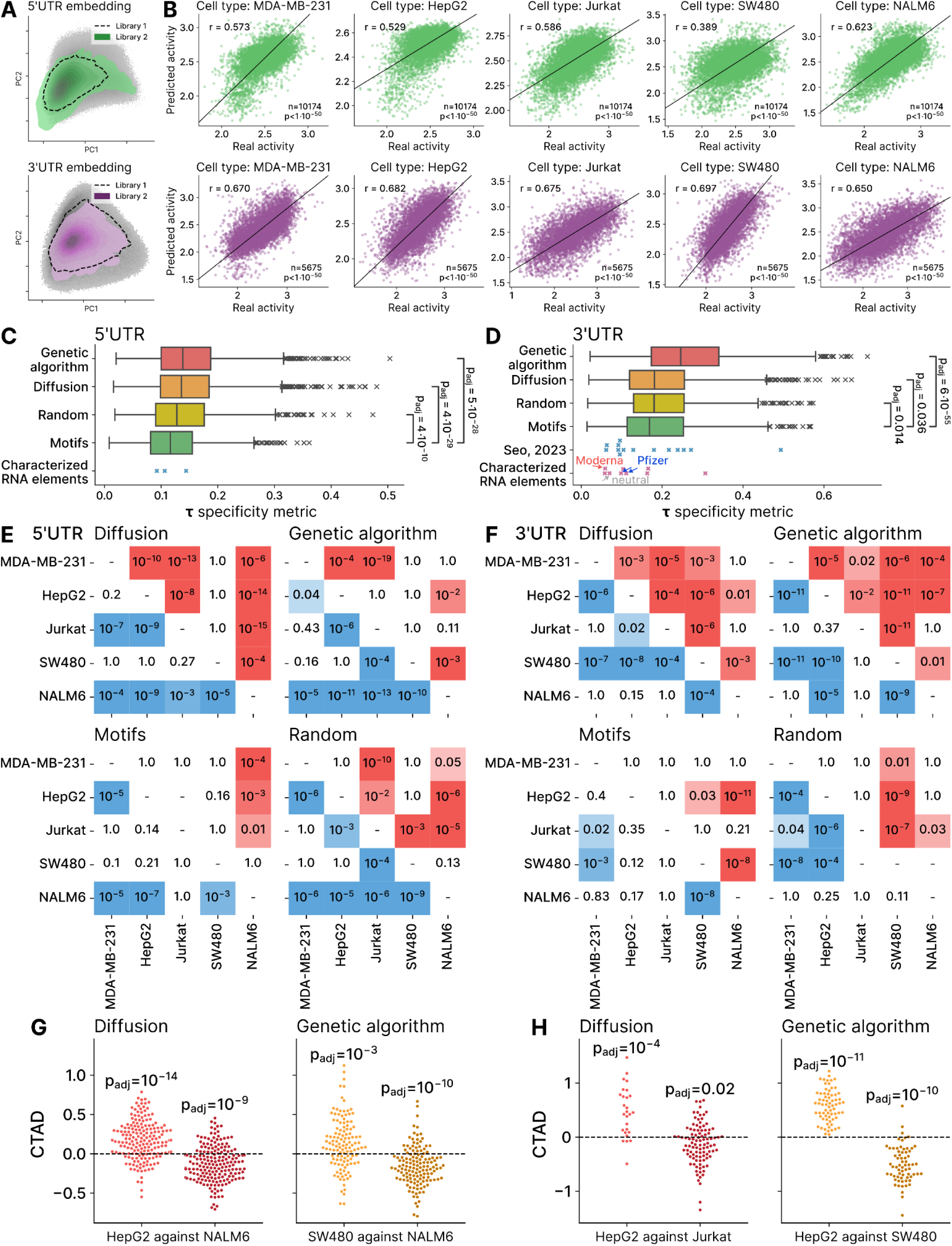
Experimental Validation Confirms Enhanced Cell Type-Specificity of UTRs Designed by the PARADE Generator. **A.** PCA of the model embeddings for Library 1 and Library 2, shown separately for 5’UTRs (top) and 3’UTRs (bottom). Random sequences are displayed in grey as the background. **B.** Scatter plots of PARADE Predictor outputs and real UTR activity experimentally measured in individual cell lines, 5’UTRs (top) and 3’UTRs (bottom). Each dot represents a single UTR sequence. **C.** Distributions of the cell type-specificity *τ* (tau) index for 5’UTRs (left) and 3’UTRs (right) across different generative methods. Box plots indicate the 10th–90th percentile range (whiskers), 25th–75th percentile range (box), and median (line). Outliers are shown as individual points. The reference RNA sequences, including 3’UTRs used in Moderna and Pfizer vaccines, are shown as a swarm plot. P-values: Mann-Whitney one-tailed nonparametric test. **D, E.** Statistical significance of the resulting cell type-specificity optimized by PARADE Generator. Each grid tile corresponds to a pairwise comparison of two target cell lines selected for CTAD (Cell Type Activity Difference) optimization. Grid tiles are colored if passing the threshold for statistical significance: red in the case of higher activity in the cell line listed in the row (positive CTAD) and blue in the case of higher activity in the column cell line (negative CTAD). **D:** 5’UTRs. **E:** 3’UTRs. P-values: Wilcoxon one-tailed nonparametric test adjusted by Holm’s multiple testing correction procedure. **F, G.** Swarm plots showing experimentally measured CTAD for optimized sequences between two target cell lines. For each plot, the CTAD values are shown for sequences optimized to favor the first cell line over the second (left) and vice versa (right). **F:** 5’UTRs. **G:** 3’UTRs. P-values: Wilcoxon one-tailed nonparametric tests comparing CTAD values with zero adjusted by Holm’s multiple testing correction procedure.

First, we assessed PARADE Predictor’s accuracy and the performance of the Diffusion method in generating sequences with the desired activity. The measured activities of Library 2 correlated well with PARADE Predictor estimates, with *r* ranging from 0.39 to 0.62 for 5’ UTR sequences and from 0.65 to 0.7 for 3’ UTR sequences (Fig. 5B), demonstrating the model’s reliability in predicting sequence activity across diverse cell lines. For sequences designed with Diffusion, the queried activity correlated with experimentally measured activity, with *r* ranging from 0.4 to 0.47 for 5’ UTRs and from 0.24 to 0.5 for 3’ UTRs (Extended Data Fig. 6A), showing that Diffusion succeeded in reaching the desired activity.

As expected, the control sequences with combinations of occurrences of activating or repressing motifs discovered with FIRE, significantly increased or decreased baseline sequence activity (p of 2⋅10^-37^ and 5⋅10^-20^ for activating motif combinations in 5’UTRs and 3’UTRs, respectively, see Extended Data Fig. 6E). These results validated both the reliability of the assay and the results of the motif discovery.

We next assessed the improvement in cell type-specificity of designed sequences included in Library 2 compared to Library 1 and the controls. The distribution of *τ* in Library 2 had a significantly higher median than in Library 1, indicating a substantial specificity improvement (*τ* of 0.084 and 0.129, for Library 1 and Library 2 respectively in 5’UTRs and 0.126 and 0.187 in 3’UTRs, see Extended Data Fig. 6F). While natural 5’ UTRs exhibited only limited cell type specificity, PARADE Generator leveraged these weak signals and achieved activity of significantly higher specificity. Furthermore, PARADE Predictor’s evaluations of *τ* correlated with those computed from experimentally measured activity for 3’ UTRs (R = 0.45), validating the model’s capacity to predict the cell type-specificity (Extended Data Fig. 6D). Further, the sequences in Library 2 demonstrated cell type-specificity on average 2x higher than Reference UTR 2 and 3.5 times higher than Reference UTR 1.

Next, we compared the performance of the generative methods with regard to the achieved cell type-specificity. Both for 5’ UTRs and 3’ UTRs, the deep learning-based methods — Genetic Algorithm, Diffusion, and Random — achieved significantly higher *τ* than Motif (adjusted p of 5*10^-28^, 4*10^-29^, 4*10^-10^ for 5’ UTRs, and 6*10^-55^, 0.036, and 0.014 for 3’ UTRs, respectively; see Fig. 5C,D). The Genetic Algorithm yielded median 2.5-fold and 4.5-fold increases in *τ* values for 3’ UTRs compared to existing RNA therapeutics, including Reference UTRs 1 and 2 (Fig. 5D), highlighting a potential area for improvement in the cell type-specificity of current mRNA vaccines. Similar trends were observed for 5’ UTRs (Fig. 4C, Extended Data Fig. 5C), although in this case, Genetic Algorithm and Diffusion yielded comparable *τ* (median values of 0.149 and 0.146, respectively). These findings illustrate the potential to significantly enhance tissue specificity in future RNA therapeutics, far surpassing that of currently available treatments (Extended Data Fig. 5F).

To further explore cell type-specificity, we grouped the sequences by the method of generation and by the target pair of cell lines for which the PARADE Generator was set to maximize CTAD. For each group, we calculated the measured activity difference between the target pair of cell types. Statistically significant differences (adjusted p<0.05) were observed for 44/80 groups of 5’ UTR sequences and for 43/80 groups of 3’ UTR sequences (Fig. 5E,F). In all cases, PARADE Generator successfully achieved the intended directionality, with activity consistently higher in the target cell line and lower in the off-target cell line (Fig. 5G,H, Extended Data Fig. 6G,H).

For 3’ UTRs, Diffusion and Genetic Algorithm yielded significant activity differences in the majority of the cell line pairs (14/20 and 13/20 groups, respectively), while the Motif and Random groups showed significant differences in 6 and 9 groups only, respectively (Fig. 5E). Similarly, for 5’ UTRs, the Diffusion and Genetic Algorithm showed significant differences in 13 and 11 out of 20 cell line pairs, respectively, while the Random group performed similarly to Diffusion (13 out of 20), and the Motif group showed significant differences in 7 out of 20 pairs only (Fig. 5D, Extended Data Fig. 5G,H). These results demonstrate that the generative algorithms were consistently successful in achieving desired cell type-specificity, especially for longer sequences such as 3’ UTRs.

### UTRs Designed by PARADE Generator Enable Cell Type-Specific Expression and Reduce Hepatotoxicity

One of the objectives of our study was to determine whether the cell type-specific RNA designs created by PARADE Generator could help alleviate therapeutic challenges, such as cargo toxicity. To test this, as our model cargo we selected CYP2E1, an enzyme whose overexpression sensitizes HepG2 cells to toxicity caused by glutathione depletion^51,52^. This cargo could be beneficial in other tissues^53^ but is harmful when expressed in the liver, and thus provides a convenient test case to demonstrate the power of cell type-specific RNA designs to minimize unintended toxic effects.

First, we tested three 3’UTR sequences with high measured CTAD between a T cell line (Jurkat) and a liver cell line (HepG2), designed by the PARADE Generator (specifically, Genetic Algorithm and Diffusion), measuring their activity individually in primary human T cells and HepG2 cell line. We synthesized FireFly luciferase-encoding mRNA (Fluc) with these UTRs in vitro, transfected the cells by electroporation, and estimated RNA activity by measuring luminescence three days later (Fig. 6A). The liver-specific UTR showed 5X higher activity in HepG2 cells, while the Jurkat-specific UTR sequences displayed 28X and 40X higher activity in T cells, respectively. This result not only demonstrates the high specificity of the individual designs but also confirms that the model trained on Jurkat cell line measurements accurately predicts activity in primary T cells.

**Figure 6.**
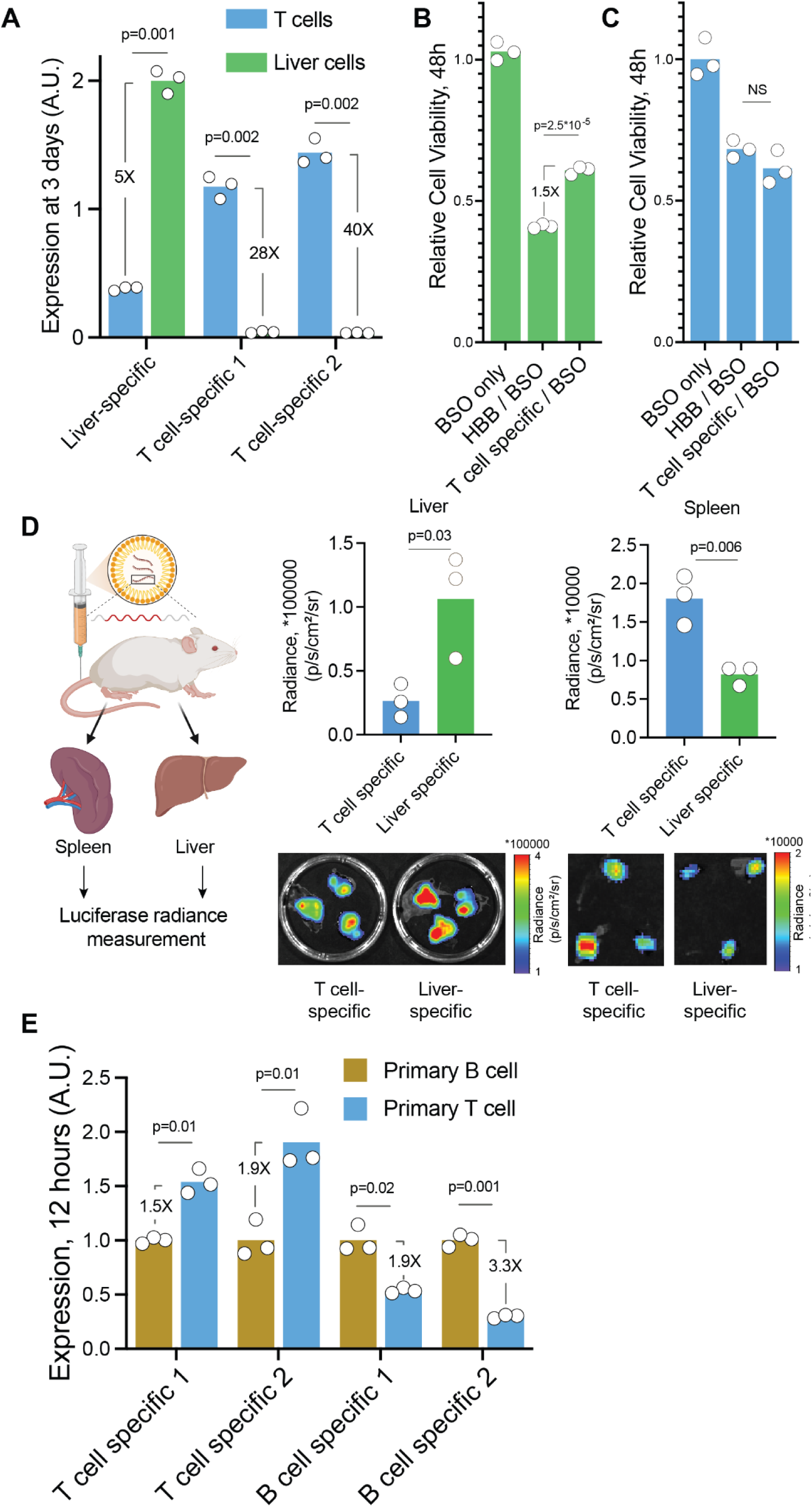
UTRs Designed by the PARADE Generator Enable Cell Type-Specific Expression and Reduce Hepatotoxicity. **A.** Luminescence of Firefly luciferase (Fluc) mRNA with HepG2-specific and Jurkat-specific UTRs in liver (HepG2) and primary human T cells, measured 3 days post-transfection. p-values: two-tailed t-test. **B.** Relative viability of HepG2 cells treated with glutathione-depleting BSO combined with CYP2E1-encoding mRNA containing either human beta-globin (HBB) or T cell-specific UTRs designed by the PARADE Generator. P-values: two-tailed t-test. **C.** Relative viability of primary human T cells treated with BSO and the same UTR configurations as in panel B. P-values: two-tailed t-test. **D.** (left) In vivo luciferase activity of Fluc mRNA with liver-specific or T cell-specific UTRs delivered via lipid nanoparticles (LNPs)^70,71^. (right, top) Radiance measurements in the liver and spleen 24 hours post-injection indicate UTR-driven specificity. (right, bottom) Representative images of spleens and livers of the treated animals. P-values: two-tailed t-test. **E.** Luminescence of Fluc mRNA with T cell- or B cell-specific UTRs in primary human T and B cells, measured 12 hours post-transfection. P-values: two-tailed t-test.

Next, to determine whether this specificity could indeed reduce hepatotoxicity, we transfected CYP2E1-encoding mRNA using either commonly used human beta-globin (HBB) 3’UTRs or the Jurkat-specific 3’UTR sequence designed by PARADE Generator. In HepG2 cells, CYP2E1 overexpression resulted in a 60% reduction in cell viability when combined with buthionine sulfoximine (BSO), a molecule that depletes glutathione. Importantly, when Jurkat-specific UTRs from the PARADE Generator were used, the cell viability increased by 48% (p = 2.5 × 10⁻⁵) compared to the HBB UTRs (Fig. 6B). Conversely, we observed no differences in the viability of T cells when using mRNAs with different UTRs, further confirming the specificity of these designs (Fig. 6C). These results highlight the therapeutic potential of novel RNA designs to safely and specifically deliver toxic payloads to target tissues, reducing the risk of off-target effects in undesired tissues, such as the liver.

Building on these results, we next evaluated the specificity of UTRs designed by PARADE Generator in animal models. We injected lipid nanoparticles containing Fluc-encoding mRNAs with either a liver-specific or a T cell-specific 3’UTR into mice (n=3 per group) via intravenous administration. After 24 hours, we isolated the livers and spleens — representing hepatocyte-enriched and T cell-enriched tissues, respectively — and measured luminescence to evaluate RNA activity (Fig. 6D). Mice injected with liver-specific UTR RNA showed significantly higher luminescence in their livers compared to the livers of mice injected with T cell-specific UTR RNA (4X difference, p=0.03), while those injected with T cell-specific UTR RNA displayed markedly higher luminescence in their spleens (2.2X difference, p=0.006). By using identical lipid nanoparticle formulation for both groups, we confirmed that these differences arose from the 3’UTR-driven RNA activity rather than variations in mRNA delivery. These findings clearly demonstrate the strong cell type-specificity of PARADE Generator designs in vivo, including their ability to drive targeted expression in distinct mouse tissues.

To further confirm that the cell type-specific activity observed in cancer cell lines reflects activity in primary human tissues, we validated the activity of four distinct 3’UTR sequences in primary human cells. Specifically, we isolated primary human T cells (CD3+) and B cells (CD3-CD19+) from peripheral blood mononuclear cells (PBMCs) and co-transfected them with Nano-luciferase-encoding mRNAs containing the 3’UTRs of interest, alongside a FireFly luciferase (Fluc) control mRNA. Twelve hours post-transfection, we measured Nano-luciferase luminescence relative to the Fluc co-transfection control. The results matched our predictions: expression differences between T and B cells ranged from 1.5X to 3.3X in the expected direction (Fig. 6E). These findings validate PARADE’s ability to generalize from cancer cell lines to primary human cells (e.g., Jurkat to T cells and Nalm6 to B cells) and demonstrate its potential to increase cell type-specificity even between closely related lineages, such as T and B cells.

### UTRs Designed by PARADE Generator Enhance mRNA Stability and Therapeutic Efficacy in Preclinical Models

Encouraged by the PARADE Generator’s success in designing cell type-specific UTRs, we sought to address another common challenge in mRNA therapeutic design: enhancing mRNA stability. We first fine-tuned PARADE Predictor on an MPRA dataset^54^ designed to assess the effect of 3’ UTRs mRNA stability, using the reporter RNA-to-DNA ratio as a proxy for mRNA stability. PARADE Predictor demonstrated superior performance compared to a simple k-mer count regression model (Pearson r of 0.58 vs. 0.43 on the held-out test subset; Fig. 7A-B).

**Figure 7.**
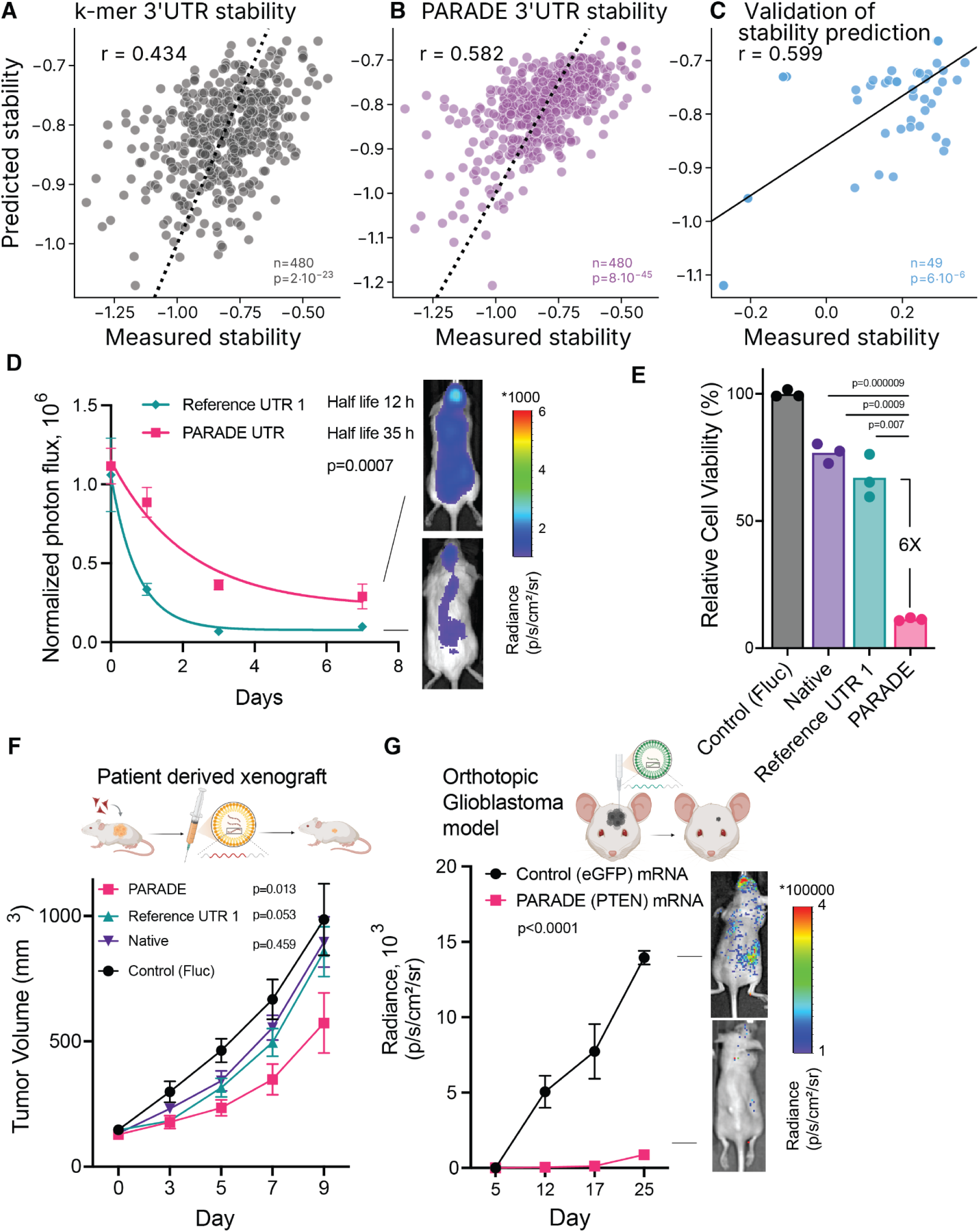
UTRs Designed by the PARADE Generator Enhance mRNA Stability and Therapeutic Efficacy in Preclinical Models. **A, B.** Comparison of eGFP reporter mRNA stability (measured as log RNA-to-DNA ratio) with k-mer regression model predictions (**A**) and PARADE Predictor outputs (**B**). The axes are shown on a logarithmic scale. **C.** Experimental validation of PARADE Predictor outputs for CD19 CAR mRNA stability, measured as the log-ratio of read counts on Day 3 to input LNPs. **D.** Longitudinal luminescence measurements of Fluc mRNA with PARADE stable UTR or Reference UTR 1 (Moderna vaccine UTR) in mice. p-values: F-test to compare one-phase decay model fits between groups. Error bars represent standard deviation, N = 4 animals. **E.** Relative viability of P16-null glioblastoma cells treated with mRNA encoding P16 with native UTR, Reference UTR 1, or PARADE stable UTR. P-values: two-tailed t-test. **F.** Tumor volume over time in patient-derived glioblastoma xenografts treated with LNPs encapsulating PTEN-encoding mRNA with PARADE stable UTR, native UTR, or Reference UTR. p-values: two-way ANOVA. Error bars represent the standard error of the mean, N = 4 animals. **G.** Radiance measurements from an orthotopic glioblastoma tumor model treated with PTEN-encoding or eGFP-encoding mRNA delivered via LNPs. P-values: two-way ANOVA. Error bars represent the standard error of the mean, N = 4 animals.

Building on these results, we experimentally validated the stability estimates from PARADE Predictor. To measure mRNA stability using a complementary approach, we synthesized RNAs encoding the therapeutically relevant protein CD19 CAR with 49 different 3’ UTRs and transfected them into primary human T cells using lipid nanoparticles (LNPs). After three days, we collected the cells, extracted RNA, and sequenced the UTRs to assess mRNA stability. The stability was quantified by calculating the ratio of RNA counts measured on Day 3 to the initial input counts. We observed a correlation of 0.60 between these measurements and the PARADE predictions (Fig. 7C), comparable to the correlation observed on a test set, thereby confirming the PARADE accuracy and generalization capabilities in modeling UTR stability.

Next, we used the PARADE Generator to design optimized UTRs aimed at increasing both mRNA stability and protein output for therapeutic applications. Specifically, we designed several 3’ UTRs, 480–600 nt in length, by combining segments of 3’ UTRs predicted to have high stability with segments predicted to have high activity in Jurkat cells. These UTRs were synthesized into FireFly luciferase-encoding mRNAs (Fluc) and transfected into primary human T cells using LNPs. Over seven days, we measured luminescence to estimate total protein production. The UTRs designed by PARADE Generator produced up to 4-fold more protein over time (area under the curve) than Reference UTR 1 and 2 (Extended Data Fig. 7A). The most effective UTR, which produced the highest total protein, was named the “PARADE stable UTR,ˮ combining a first half optimized for high stability with a second half optimized for high activity (Extended Data Fig. 7B).

To test the impact of these findings in animals, we evaluated the stability of Fluc mRNA with either Reference UTR 1 or the PARADE stable UTR in mice. We intravenously injected LNPs containing these mRNAs and measured luminescence over a week (Fig. 7D). On Day 7, the PARADE stable UTR group of animals showed 3X higher luminescence than the Reference UTR 1 group, with an mRNA half-life 3X longer as well (35 hours vs. 12 hours). These results demonstrated that UTRs created by PARADE Generator not only improved mRNA stability but also enhanced protein production in animal models.

With this robust stability and activity, we explored how UTRs designed by PARADE Generator could improve the expression of therapeutic proteins across multiple cancer models. Since durable expression of oncosuppressor proteins like PTEN, P16, or P53 leads to cell death in cancer cells with null mutations in these genes^55,56^, we used these experimental cancer systems to link mRNA stability to measurable phenotypic outcomes. First, we tested this in P16-null H4 neuroglioma cells by transfecting P16 mRNAs with native UTRs, Reference UTR 1, or PARADE stable UTR. While the native mRNA caused only a 30% reduction in cell viability, the PARADE stable UTR improved the effect significantly, reducing viability 6X more than Reference UTR 1 (Fig. 7E).

To confirm these effects in vivo, we used an entirely independent patient-derived xenograft model of glioblastoma in mice. We administered P16 mRNA via LNPs containing native, Reference UTR 1, or PARADE stable UTR to mice intratumorally (5 injections of 2 mg/kg mRNA every 2 days, N=12 animals per group) and monitored the tumor growth. By Day 9, the PARADE stable UTR group exhibited tumors that were 33% smaller than those in the Reference UTR 1 group and 36% smaller than those in the native mRNA group (Fig. 7F). RT-qPCR quantification of biopsies on Day 10 revealed 22-fold higher mRNA abundance in tumors treated with PARADE stable UTR RNA compared to Reference UTR RNA, confirming that improved mRNA stability drove these therapeutic effects (Extended Data Fig. 7C).

Finally, we extended our evaluation to an orthotopic mouse model of neuroglioma. In this model, we established intracranial glioblastomas by injecting H4 cells into mice, and delivered PTEN-encoding mRNA directly into the tumors using convection-enhanced delivery (CED). Glioblastomas are highly aggressive and lack effective treatments, making this a critical test of PARADE Generator’s potential. Administering PTEN mRNA with the PARADE stable UTR significantly reduced tumor growth (2 injections of 0.03 ug mRNA every 7 days, N=4 animals per group, p = 0.0002) and improved survival (p = 0.034) compared to the control GFP mRNA (Fig. 7G, Extended Data Fig. 7D). These findings highlight PARADE Generator’s ability to enhance mRNA stability across multiple independent cargos, i.e. luciferase, P16, and PTEN, with significant phenotypic consequences in preclinical models of hard-to-treat diseases, such as glioblastoma.

Taken together, we have established PARADE as a versatile and powerful framework for designing UTRs with tailored activity, cell type-specificity, and enhanced stability. Using high-throughput MPRA data, PARADE Predictor accurately evaluated regulatory element effects, identified key motifs driving cell-specific activity, and guided the generation of optimized UTR sequences. Experimental validation across diverse models — cell lines, primary cells, and animal models — demonstrated PARADE’s ability to mitigate cargo toxicity, enhance protein production, and improve therapeutic efficacy. These findings highlight the broad potential of UTRs designed with PARADE Generator for advancing RNA-based therapies and pave the way for further exploration of its applications in clinical research.

## Discussion

In this study, we utilized generative AI to design UTRs that enhance cell type specificity and stability, key factors in improving the therapeutic effectiveness of mRNA. By leveraging large-scale MPRA data, our PARADE Framework integrates predictive modeling and generative methods to create UTRs that outperform naturally occurring sequences and those of existing RNA therapeutics in terms of cell type-specificity. The generative approaches employed within the PARADE Generator, such as Diffusion and Genetic Algorithm, were particularly effective in designing UTRs with tailored activity, as detailed in the subsequent discussion of their comparative performance. These findings highlight PARADE Framework’s dual capability to predict RNA activity with high accuracy and to generate novel UTR sequences capable of addressing key challenges in mRNA-based therapeutics, such as off-target effects and tissue toxicity.

### Generative Models Enhance Specificity Over Random and Natural Sequences

Our results show that generative AI methods — particularly Diffusion and Genetic Algorithm — achieve significantly higher cell type-specificity for longer sequences, such as 240 nt-long 3’ UTRs, than random sampling or motif seeding techniques. This finding aligns with the hypothesis that AI-driven models are more effective at exploring the complex multidimensional sequence space, especially where longer sequences allow for non-linear interactions between individual regulatory elements. For shorter sequences, such as 50 nt-long 5’ UTRs, the difference between the generative methods and random sampling was less pronounced, likely due to the limited scope for motif interactions in shorter regions. Yet, even for 5’ UTR it became possible to achieve non-uniform activity across cell types, highlighting the cell type-specific regulatory layer that is often overlooked^57^.

These insights emphasize a broader challenge in RNA therapeutics: naturally occurring RNAs, shaped by evolutionary pressure, do not reach extreme levels of stability or specificity due to the need for flexible regulation. One explanation is that physiological RNAs are depleted of strong regulatory signals such as high-affinity RBP binding sites, leading to promiscuous and weak RBP binding as demonstrated for C5^58^. In theory, the specificity of RNA activity could be enhanced by learning the functional motifs from natural RNAs, strengthening them, and arranging them into novel combinations not present in native transcripts. Deep learning models are specifically useful in this context, as they account for complex non-linear interactions of both known and not yet studied regulatory elements^59^. As demonstrated in Fig. 5C, the sequences created by PARADE Generator show significantly higher cell type-specificity than naturally occurring RNAs, underscoring the potential of generative models to overcome evolutionary constraints. While previous efforts have focused solely on designing UTRs to improve stability and translation efficiency using AI-based models^14,22,60–62^ — including applications leveraging AI to enhance mRNA stability^63^ — our work extends these efforts by also addressing cell type-specificity, offering a more comprehensive and flexible solution for mRNA therapeutic design.

### Implications for RNA Therapeutics

The PARADE Generator’s ability to design RNA sequences with enhanced tissue specificity and stability offers significant potential for advancing mRNA therapeutics. By tailoring these properties, the PARADE Framework can address key challenges in therapeutic mRNA design, such as reducing off-target effects and improving the durability of the RNA treatment. The combination of these attributes in future RNA designs could enable highly targeted therapies with increased efficacy and safety.

Looking ahead, the PARADE Generator could be applied to develop mRNA molecules that are both highly stable and tissue-specific. This would open up new therapeutic possibilities, such as targeting heart tissue with a growth factor mRNA to promote recovery after a heart attack without causing unwanted effects in other tissues^64^. Similarly, more precise targeting of therapeutic mRNA for neurodegenerative diseases could reduce adverse effects in non-target areas of the brain^65^.

### Current Limitations and Future Directions

While PARADE succeeded in quantitative predictions of UTR activity, the median correlation of 0.82 does not yet match the correlation of 0.95 observed in yeast promoter-interrogating MPRAs ^66^. Further, the accuracy of the activity measurements depends on data preprocessing (such as normalization) and the plasmid design ^29^. Particularly, UTRs that fall into the first cell sorting bin may be subject to transcriptional noise^67^, including noise from cryptic IRES promoters^68^. Expanding the dataset to include a broader range of cell types and biological contexts could boost predictive correlations achieved by PARADE and further enhance its overall performance. Incorporating multiple independent cell lines for each cell type, rather than relying on a single cell line per tissue, will improve the robustness and generalizability of the identified UTRs across biological systems. By combining PARADE Framework with iterative active learning approaches, the framework can be continuously refined, pushing the boundaries of specificity and therapeutic efficacy.

While the PARADE Framework has demonstrated success in generating UTRs that are both more stable and more tissue-specific, extending this capability to full-length mRNA design, including both the untranslated and coding regions, remains a key future direction as the coding region also plays a crucial role in determining the cell type-specificity of the post-transcriptional gene expression control^69^. Generative models capable of designing entire mRNAs on top of the PARADE Framework presented here could further enhance the precision and efficacy of RNA therapeutics.

In summary, the demonstrated ability of PARADE Framework to design RNAs with tailored properties — such as increased tissue specificity and improved stability — marks a significant advancement in RNA-based therapeutics. As these methods are further refined, the potential to address unmet medical needs with safe, effective, and precisely targeted mRNA therapies will continue to expand.

## Extended Figures

**Extended Data Figure 1.**
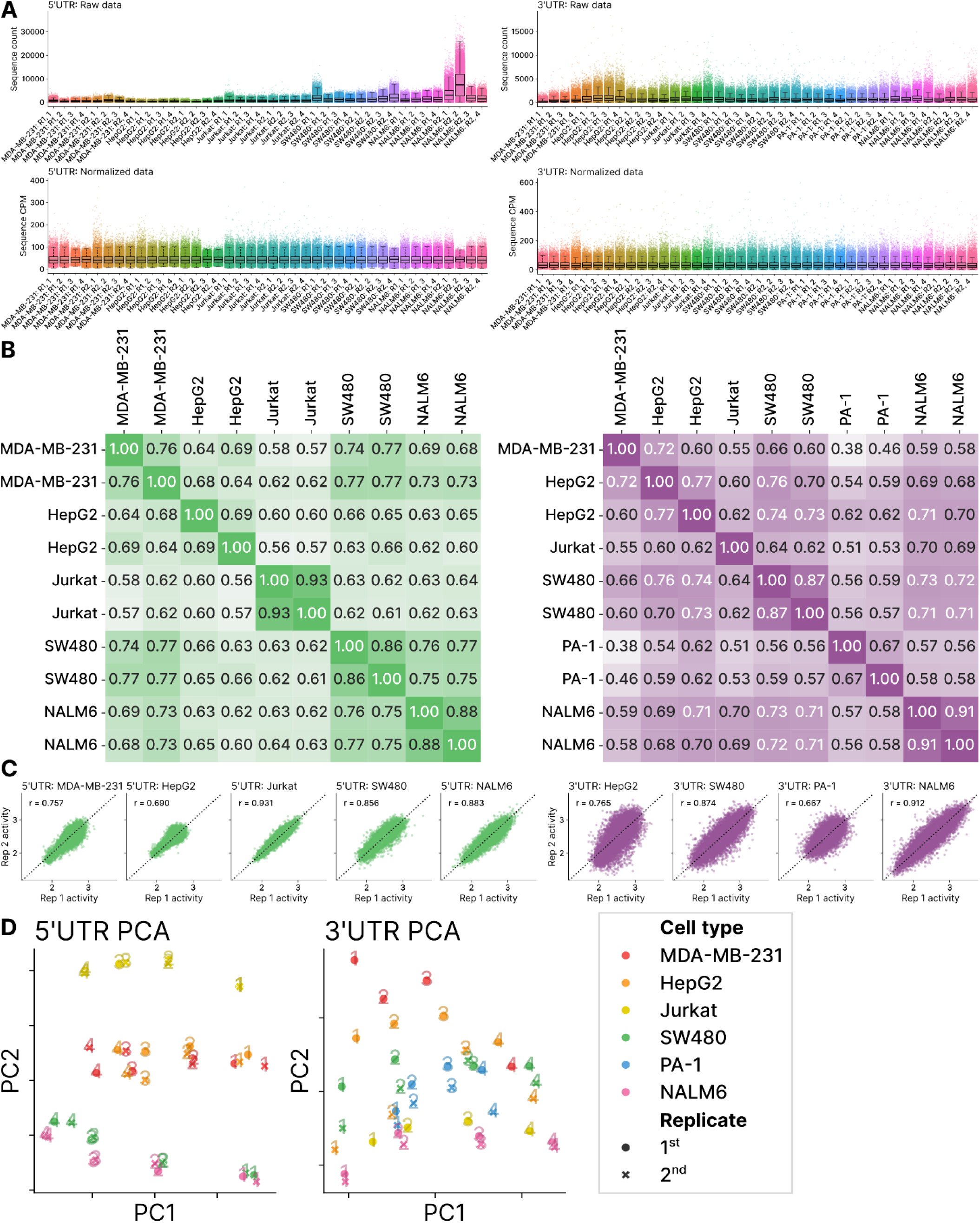
Experimental Metrics Demonstrating Robustness and Consistency of MPRA Assay Across Cell Lines for Library 1. **A.** Sequence count distributions for 5’UTRs (left) and 3’UTRs (right) across all individual libraries, before (top) and after normalization (bottom) for different cell lines. Each boxplot represents data for a single library. **B.** Pairwise Pearson correlation heatmaps showing reproducibility of sequence activity between replicates and across the cell lines for 5’UTRs (left) and 3’UTRs (right). Color scale saturation: Pearson correlation value. **C.** Scatterplots comparing sequence activity measurements between replicates for each cell line. Pearson correlation coefficients (r) are shown on the plots for 5’UTRs (green) and 3’UTRs (purple). **D.** PCA of 5’UTR (left) and 3’UTR (right) libraries, points are colored by the cell line. The markers are numbered according to the sorting bin numbers and shaped according to replicates.

**Extended Data Figure 2.**
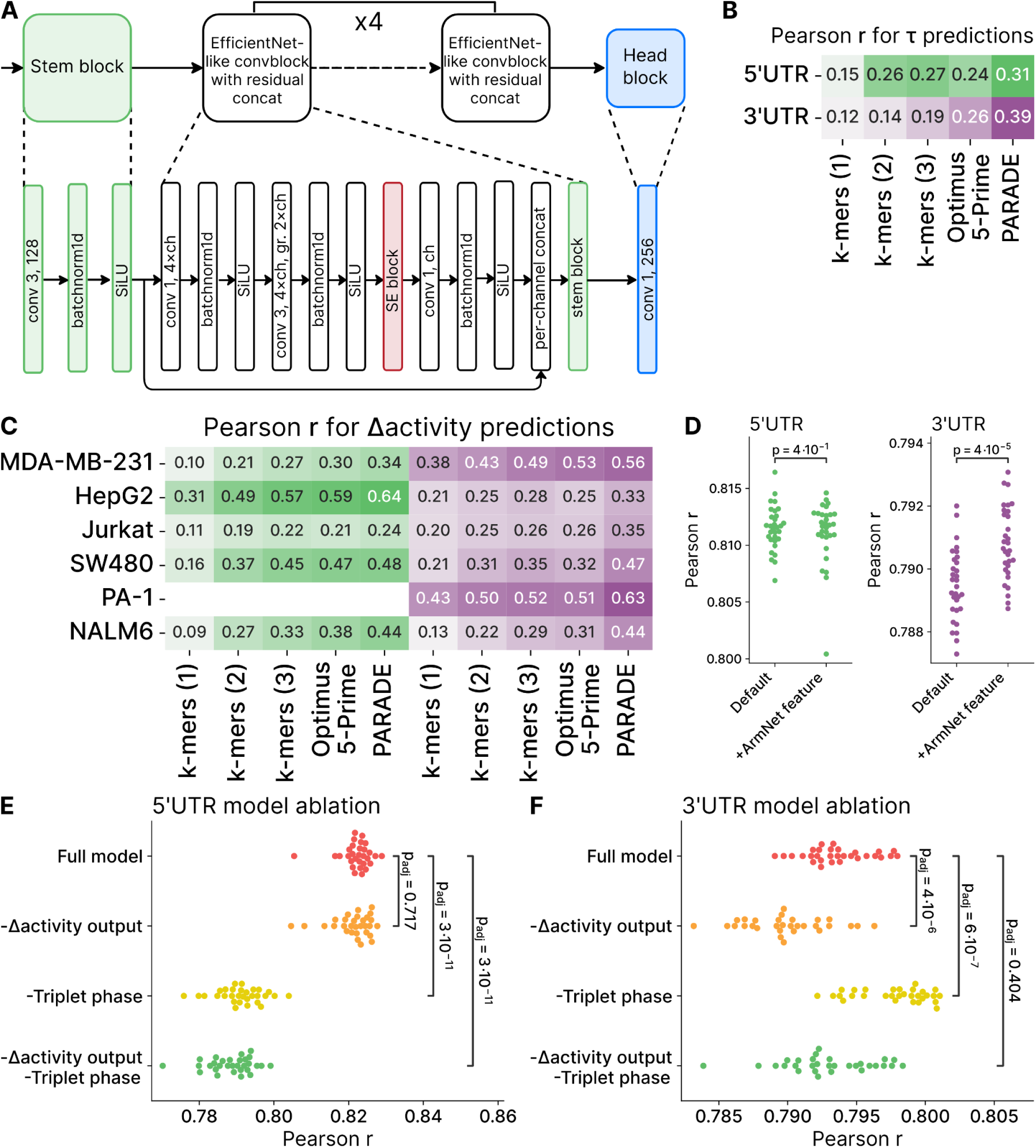
Performance Metrics for PARADE Predictor’s Evaluation of UTR Activity and Specificity. **A.** Schematic of the PARADE Predictor architecture. **B.** Pearson correlation (r) for *τ* specificity metric calculated using predicted and measured UTR activity values for 5’UTRs (green) and 3’UTRs (purple) across PARADE Predictor and baseline models (k-mer and Optimus-5-prime). **C.** Pearson correlation (r) values for activity deviation Δ predictions for PARADE Predictor and baseline models across different cell lines, shown separately for 5’UTRs (green) and 3’UTRs (purple). **D.** Impact of using additional RNA structure features predicted by ArmNet^72^ on model performance. Pearson correlation of predicted and measured values for 5’UTR (green) and 3’UTR (purple) are shown. P-values: Mann-Whitney nonparametric test. **E, F.** Ablation studies of PARADE Predictor 5’UTR (**E**) and 3’UTR (**F**) models. The Pearson correlation for activity predictions is shown for the full model and the models with specific features removed (Δ activity output, triplet phase, or both). P-values: Mann-Whitney nonparametric test.

**Extended Data Figure 3.**
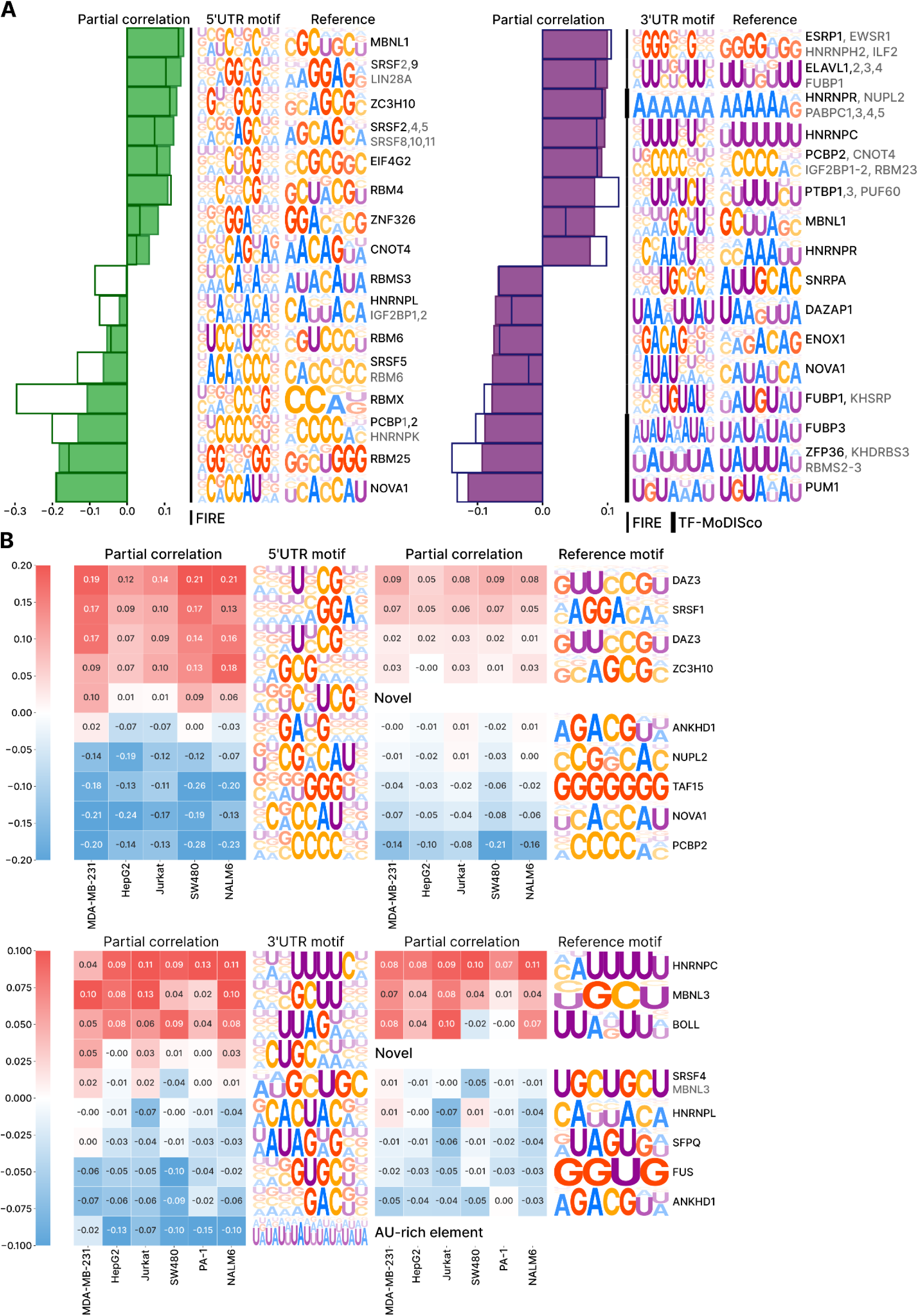
Expanded Identification of Cell Type-Specific Motifs and Their Regulatory Annotations. **A.** Waterfall plots showing 9+9 motifs with the highest and the lowest mean partial correlation between motif scores and experimentally measured sequence activity for 5’UTRs (green, left) and 3’UTRs (purple, right). RNA-binding protein motifs were identified using FIRE and TF-MoDISco. Filled bars indicate the mean partial correlation (corrected for the nucleotide composition) of the motif scores and the measured activity. Colored bars: discovered motifs, empty bars: the respective reference motifs. **B.** Heatmaps of partial correlations for select motifs demonstrating cell type-specific regulatory effects across six tested cell lines. Left: 5’UTR motifs; right: 3’UTR motifs.

**Extended Data Figure 4.**
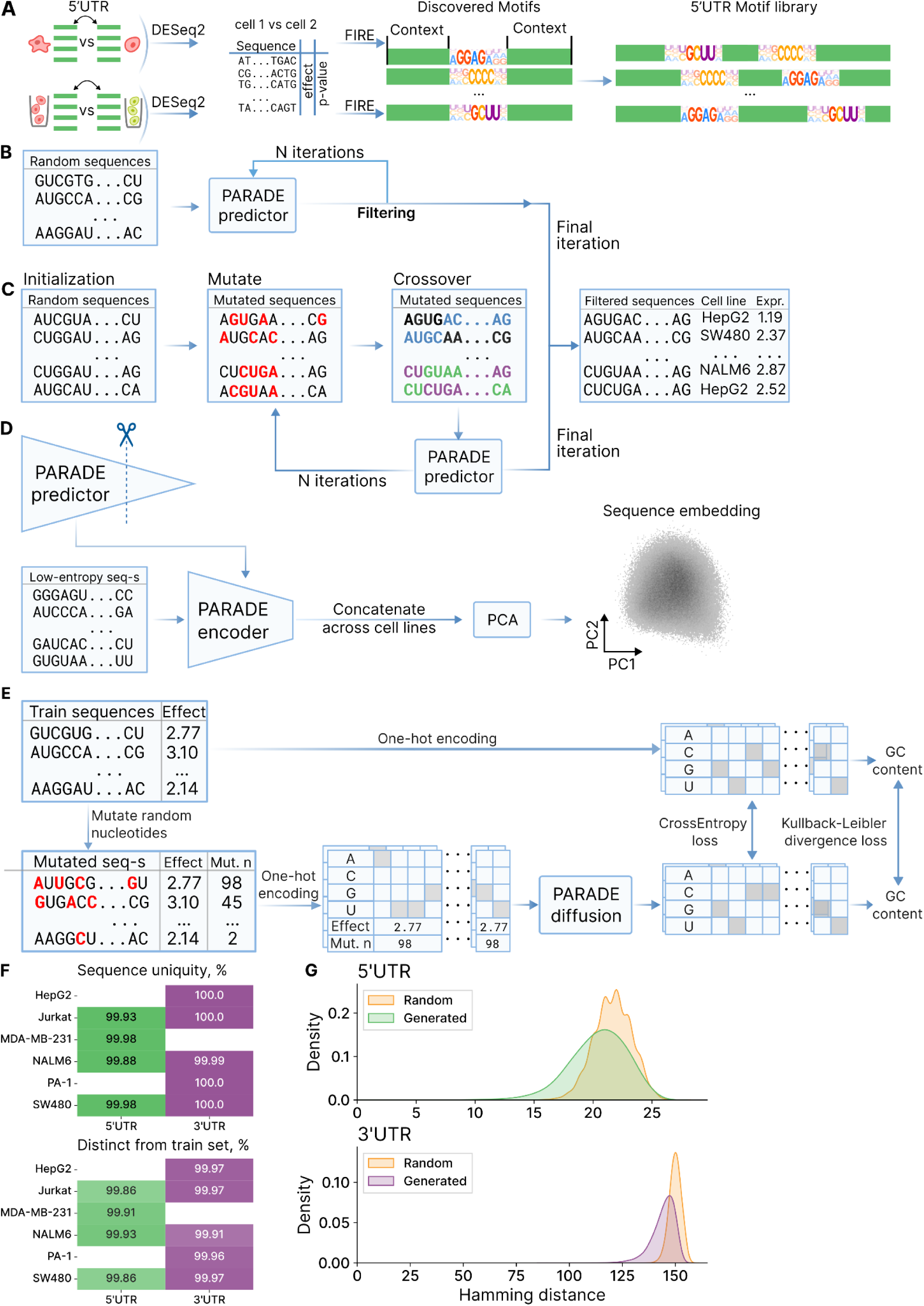
Overview of Generative Algorithms for Cell Type-Specific UTR Design. **A.** Workflow for motif discovery from UTRs differentially active in two cell types, followed by the generation of motif-based UTR libraries. **B.** Diffusion generative approach, where random sequences undergo iterative refinement using the PARADE Predictor to optimize for desired activity. **C.** Genetic Algorithm workflow, involving initialization with random sequences, iterative mutation and crossover, and final selection of optimized sequences using the PARADE Predictor. **D.** Latent space exploration: PCA of concatenated sequence embeddings generated by the PARADE Predictor across cell lines. **E.** Proportion of Diffusion-generated 5’UTR and 3’UTR sequences that are unique (top) or distinct from training data (bottom) across cell line-specific models. **F.** Kernel density estimates of Hamming distances between generated or random sequences and their closest counterparts in the training set (Library 1) for 5’UTRs (top) and 3’UTRs (bottom). Lower values indicate greater similarity to the sequences of the training set.

**Extended Data Figure 5.**
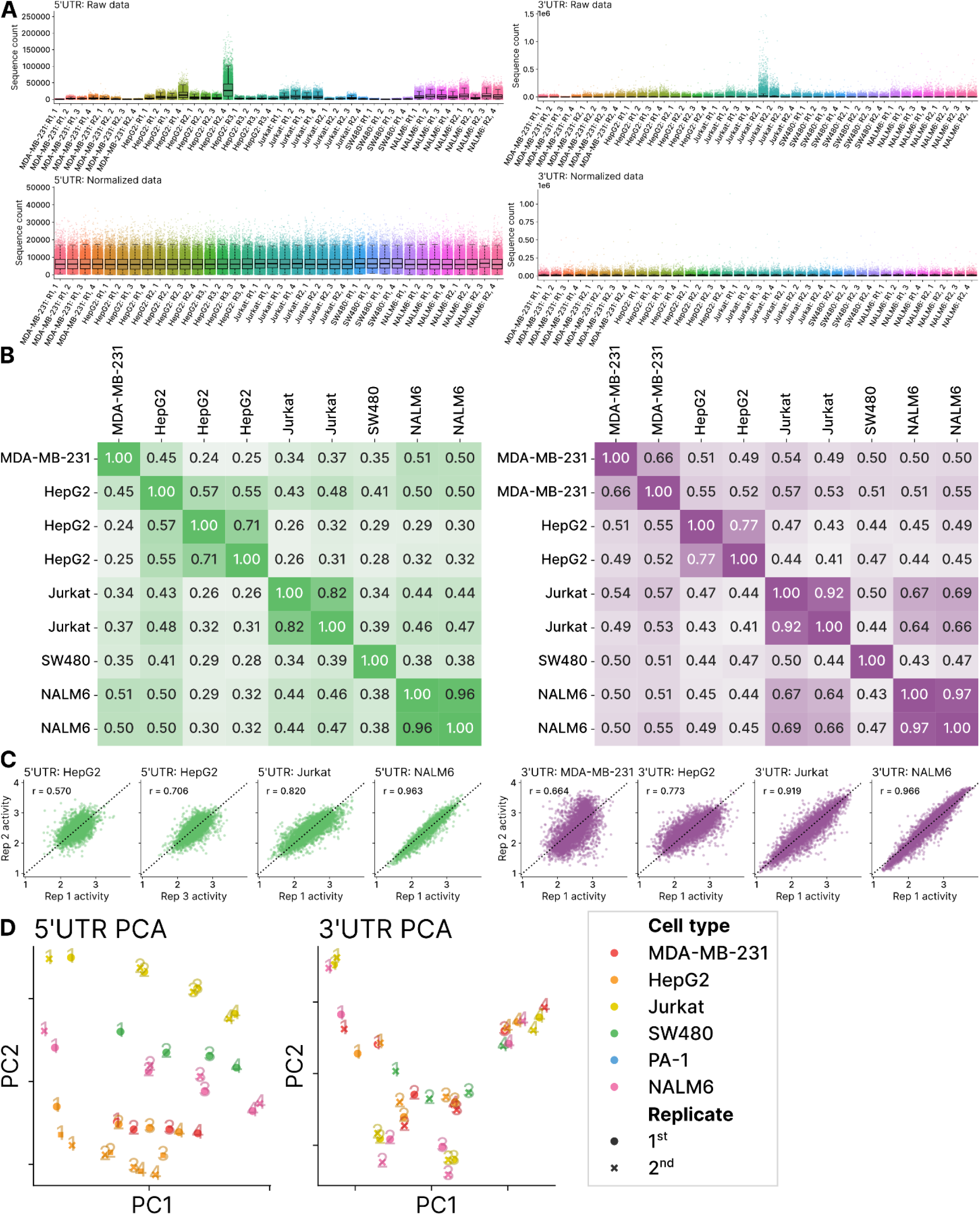
Metrics Demonstrating Robustness and Consistency of MPRA Assay Across Cell Lines for Library 2. **A.** Sequence count distributions for 5’UTRs (left) and 3’UTRs (right) across all individual libraries for Library 2, shown before (top) and after normalization (bottom) across six cell lines. Each boxplot represents data from a single library. **B.** Pairwise Pearson correlation heatmaps for Library 2, illustrating the reproducibility of sequence activity measurements between replicates and across cell lines for 5’UTRs (left) and 3’UTRs (right). Color scale saturation: Pearson correlation. **C.** Scatter plots comparing sequence activity measurements between replicates for each cell line in Library 2, with Pearson correlation coefficients (r) shown for 5’UTRs (green) and 3’UTRs (purple). **D.** PCA of 5’UTR (left) and 3’UTR (right) libraries in Library 2, colored by cell line. Markers are numbered to reflect sorting bin numbers and shaped to indicate the replicate identity.

**Extended Data Figure 6.**
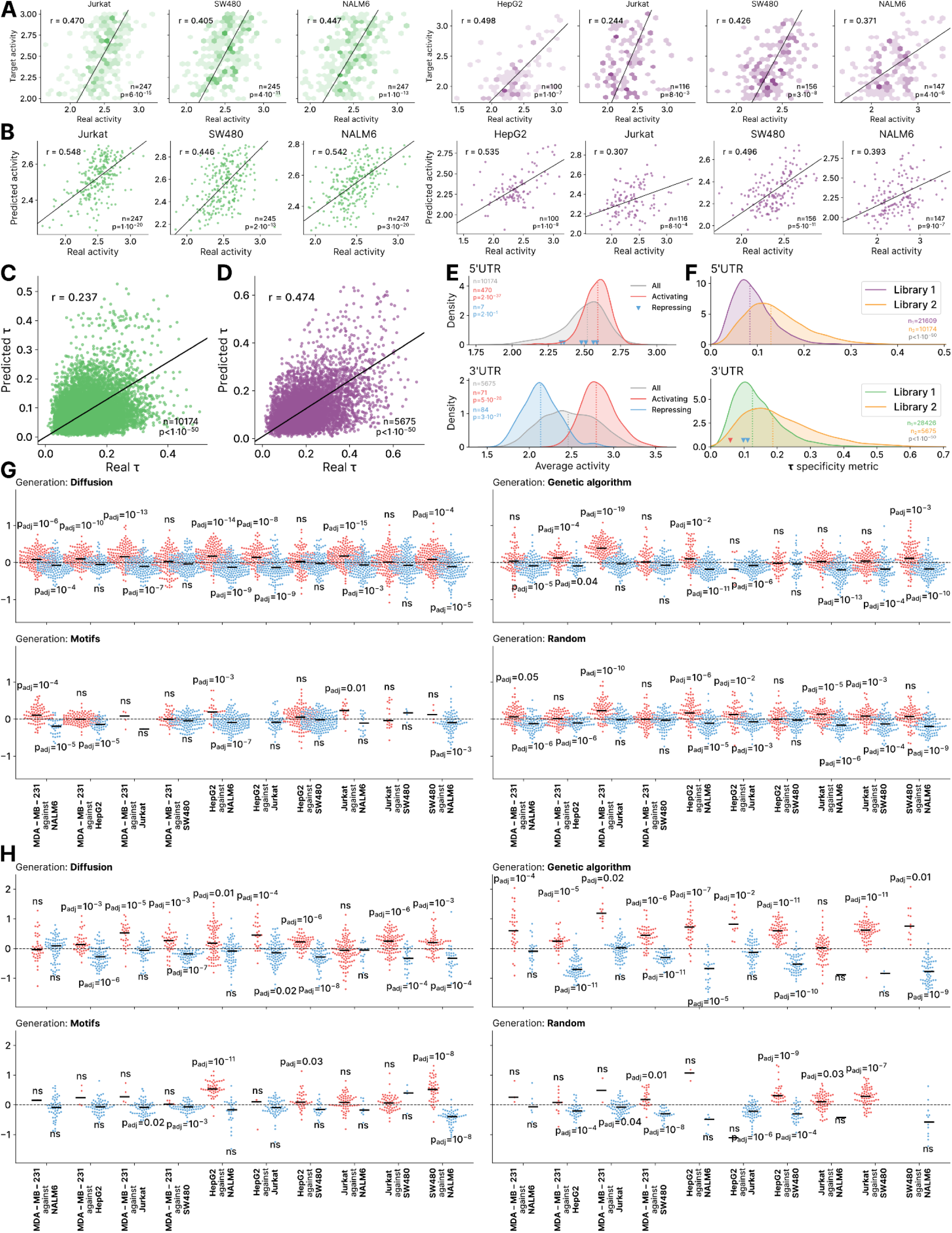
Performance of PARADE Framework in Designing and Predicting UTR Activity and Specificity. **A, B.** The correlation between experimentally measured activity and the target activity intended during PARADE Generation sequence design (**A**) or the PARADE Predictor outputs (**B**) is shown for Diffusion-generated sequences. Pearson correlation coefficients (r) are indicated on the plots; the results are displayed for 5’UTRs (left) and 3’UTRs (right). **C, D.** Comparison of the *τ* cell type-specificity index computed from predicted and measured activity values for 5’UTRs (**C**) and 3’UTRs (**D**) in Library 2. Pearson correlation coefficients (r) and p-values are indicated. **E.** Sequence activity distributions in Library 2, kernel density estimation. Distributions are shown for sequences designed with activating or repressing motifs only, separately for 5’UTRs (top) and 3’UTRs (bottom). Medians are shown with dashed lines. P-values: one-tailed Mann-Whitney U test. **F.** Comparison of the *τ* specificity distributions for 5’UTRs (top) and 3’UTRs (bottom) for Library 1 and Library 2, demonstrating increased cell type-specificity of Library 2 sequences. Reference sequences are indicated as follows: Reference UTR 1 (Moderna COVID vaccine UTR, red), Reference UTR 2 (Pfizer COVID vaccine UTR, blue). Medians are shown with dashed lines. P-values: one-tailed Mann-Whitney U test. **G, H.** Swarm plots of Cell Type Activity Difference (CTAD) measurements for optimized sequences targeting only one of two selected cell lines. For each plot, CTAD values are shown for sequences optimized to favor the first cell line over the second (left, red) and the second over the first (right, blue). Results are grouped by the sequence generation method. Individual dots represent individual sequences. Data are shown for 5’UTRs (**G**) and 3’UTRs (**H**). Boxplots represent the median (center line), interquartile range (box edges: 25th and 75th percentiles), and the whiskers (10th to 90th percentiles). P-values: one-tailed Wilcoxon tests with Holm’s correction.

**Extended Data Figure 7.**
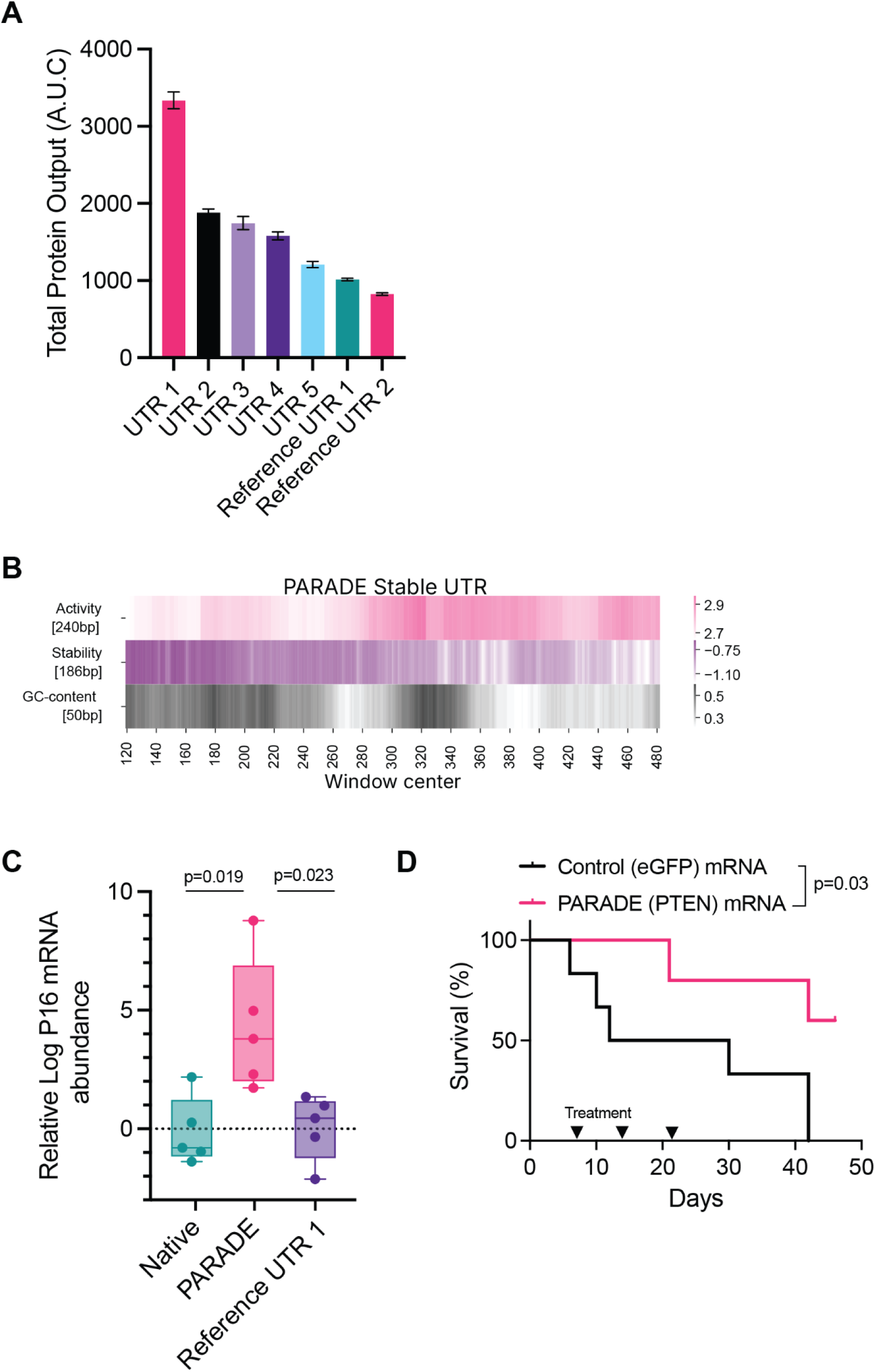
UTRs Designed by the PARADE Generator Enhance RNA Stability and Therapeutic Efficacy in Preclinical Models. **A.** Total protein output from mRNAs carrying UTRs designed by PARADE Generator compared with Reference Optimized UTRs 1 and 2 (COVID vaccine UTRs), calculated as the area under the curve (AUC). Error bars represent the standard error of the mean (SEM) from three replicates. P-values: two-tailed t-test. **B.** Sliding window average predictions of Activity and Stability for the PARADE Stable UTR **C.** Relative P16 mRNA abundance in tumor biopsies collected on day 10 from mice treated with mRNA containing native UTR, Reference UTR 1, or PARADE stable UTR, measured by RT-qPCR. Error bars represent SEM. P-values: two-tailed t-test. **D.** Kaplan-Meier survival analysis of mice treated with lipid nanoparticles (LNPs) encapsulating PTEN mRNA with the PARADE stable UTR versus eGFP control mRNA in an orthotopic glioblastoma model. Statistical significance was determined using a log-rank test.

## Methods

### Cell culture

All cells were cultured in a 37°C 5% CO_2_ humidified incubator. The cell lines Jurkat, Nalm-6, SW-480, PA-1, MDA-MB-231, and HepG2 were purchased from ATCC (ATCC identifiers tib-152, crl-3273, ccl-228, crl-1572, htb-26 and hb-8065) and were cultured in RPMI-1640 medium supplemented with 10% FBS, glucose (2 g/L), L-glutamine (2 mM), 25 mM HEPES, penicillin (100 units/mL), streptomycin (100 μg/mL) and amphotericin B (1 μg/mL) (Gibco). All cell lines were routinely screened for mycoplasma with a PCR-based assay.

### Reporter vector library construction and cloning

To construct the library, the Translation Efficiency (TE) values were compared between the Ribosome Profiling datasets collected in naive mouse CD4+ T cells^26,27^ and mouse livers^25^ at the gene level, using DESeq2^73^. Genes with significant changes were selected (Log_2_ Fold Change > 1, negative log_10_ of adjusted P-value > 10). The 5′UTRs and 3′UTRs of human orthologs of these genes and all other genes in the human genome were split into overlapping chunks of 50 nt (for 5′UTRs) and 240 nt (for 3′UTRs), respectively. From these, 6,000 fragments of UTRs were randomly selected from human orthologs of genes with significant changes in TE values, 14,000 fragments were randomly selected from other human genes (one per gene), and 10,000 fragments were randomly selected from all genes in the human genome.

The selected 5′UTR and 3′UTR sequences were cloned as described previously by ^54,74^. For 5′UTRs, the sequences were flanked with the adapters TCCCTTGGAGAACCACCTTGGGTCTCnCGTC-AGATC at the 5′ end and GCCACC-ATGGnGAGACCTAAGCTGGAAACAGCATAGCAAG at the 3′ end. For 3′UTRs, the sequences were flanked with the adapters TCCCTTGGAGAACCACCTTGACGCGT at the 5′ end and TTAATTAATAAGCTGGAAACAGCATAGCAAG at the 3′ end. PCR adapters, restriction sites, and other sequences were added based on prior recommendations.

The oligonucleotide library was synthesized commercially and amplified with minimal PCR cycles as per manufacturer protocols. Amplified libraries were digested with MluI and PacI for 3′UTRs, or BsaI for 5′UTRs (NEB). The cloning vectors used were pMK053 (Addgene #213963) for 3′UTRs, which was digested with MluI, PacI, and rSAP (NEB), and pMK089 (Addgene #213964) for 5′UTRs, which was digested with BspMI and rSAP (NEB). Digested DNA was purified using the DNA Clean & Concentrator-5 kit (Zymo Research) and polyacrylamide gel electrophoresis, with gel percentages adjusted based on fragment size.

Ligations were performed using T4 DNA ligase (NEB), and the ligation products were transformed into Endura Competent Cells (Lucigen) via electroporation for larger libraries. Chemical transformation was used for smaller libraries. The transformed libraries were PCR-amplified and sequenced on a MiSeq Illumina sequencer to confirm the diversity and correctness of the library.

### Massively Parallel Reporter Assays

The MPRA assays were performed as described in^54^. The DNA library was co-transfected with pCMV-dR8.91 and pMD2.G plasmids using TransIT-Lenti (Mirus) into HEK293 cells, following the manufacturer’s protocol. The virus was harvested 48 hours post-transfection and passed through a 0.45 µm filter. The cell lines Jurkat, Nalm-6, SW-480, PA-1, MDA-MB-231, and HepG2 were then transduced overnight with the filtered virus in the presence of 8 µg/mL polybrene (Millipore); the amount of virus used was optimized to ensure an infection rate of 30%. The infected cells were selected with 2 µg/mL puromycin (Gibco). Cells were harvested for sorting and analysis on a BD FACSaria II sorter. The distribution of GFP to mCherry ratios was calculated for sorting the library into subpopulations: we gated the population into 4 bins each containing 25% of the total number of cells. A total of 5 million cells were collected in each bin to ensure sufficient representation of sequences in the population in two replicates each. For each subpopulation, genomic DNA was extracted with NucleoSpin Blood L Vacuum Genomic DNA (Macherey Nagel).

Genomic DNA (gDNA) was amplified by a 24-cycle PCR reaction using Q5 polymerase (NEB). For 3’UTR libraries, the forward primer was designed as: AATGATACGGCGACCACCGAGATCTACAC | NNNNNNNN | ACACTCTTTCCCTACACGACGCTCTTCCGATCT | XXX | GTGGTCTGGATCCACCGGTCC. This primer includes, in order, the Illumina P5 adapter, the i5 index (NNNNNNNN, where N represents a sequence of any nucleotide), the TruSeq Read 1 primer site, a staggering region (XXX, a mix of 4 primers with 1 to 4 random nucleotides to increase library diversity), and the primer binding site located downstream of the insert. The reverse primer was designed as: CAAGCAGAAGACGGCATACGAGAT | NNNNNNNN | GTGACTGGAGTTCAGACGTGTGCTCTTCCGATC | XXX | ACTGCTAGCTAGATGACTAAACGCGT. This primer includes, in order, the Illumina P7 adapter, the i7 index (NNNNNNNN, where N represents a sequence of any nucleotide), the TruSeq Read 2 primer site, a staggering region (XXX, a mix of 4 primers with 1 to 4 random nucleotides to increase library diversity), and the primer binding site located upstream of the insert.

For 5’UTR libraries, the forward primer was designed as: AATGATACGGCGACCACCGAGATCTACAC | NNNNNNNN | ACACTCTTTCCCTACACGACGCTCTTCCGATCT | XXX | GAGCTCGTTTAGTGAACCGTCAGATC. This primer includes, in order, the Illumina P5 adapter, the i5 index, the TruSeq Read 1 primer site, a staggering region, and the primer binding site downstream of the insert. The reverse primer was designed as: CAAGCAGAAGACGGCATACGAGAT | NNNNNNNN | GTGACTGGAGTTCAGACGTGTGCTCTTCCGATC | XXX | CCGGTGGTGCAGATGAACTTC. This primer includes, in order, the Illumina P7 adapter, the i7 index, the TruSeq Read 2 primer site, a staggering region, and the primer binding site located upstream of the insert.

Different i7 indices were used for different bins to allow multiplexing, with different i5 indices applied across the two replicates. The amplified DNA libraries were size-purified using the Select-a-Size DNA Clean & Concentrator MagBead Kit (Zymo). Deep sequencing was then performed on a NovaSeq X platform (Illumina) at the UCSF Center for Advanced Technologies.

The adapter sequences were removed using Cutadapt^75^. For RNA libraries, UMIs were then removed from the reads and appended to read names using UMI tools^76^. The reads were matched to the UTRs using BWA-MEM ^77^. The read counts were obtained using featureCounts^78^.

### MPRA data normalization

We implemented an advanced normalization to unify the read counts distribution across libraries. To this end, we selected a common model distribution (zero-inflated negative binomial, ZINB, for 5’UTRs and mixture of two ZINBs for 3’UTRs) and employed PyMC3 to fit the observed data to the model for each sequencing batch (i.e. bin-replicate pair). For fitting, uniform priors were used for ѱ (probability of zero) and n (number of trials), and the Beta prior was used for p (the success probability). For 5’UTRs and 3’UTRs, the bin-replicate pair with the average read count closest to the global mean was chosen as the reference. After obtaining the fitted distributions for each sequencing batch, we performed a quantile normalization of values with regard to the reference distribution. The UTR activity was then estimated as the weighted average of the bin numbers (1-4) with the sum of normalized counts in the bin in all replicates as the bin weights. The same normalization procedure was applied to the sequence counts of Library 2 and to the stability MRPA data, where RNA and DNA counts were normalized separately.

### PARADE Predictor

To create a predictive model, we modified the LegNet architecture^37,66^ in order to adapt it to a smaller volume of training data (see Extended Data Fig. 2A). The following modifications were applied: the amount of LegNet blocks was reduced to 5 of sizes 128 (stem), 64, 64, 32, 32, followed by a single linear layer with 2 outputs. The model was trained to minimize the mean square error (MSE Loss) for 10 epochs with an AdamW optimizer (weight_decay=0.1), and OneCycle learning rate scheduler (max_lr=0.01). The batch size was fixed at 1024 sequences per batch.

### RNA structure features

UTRs can influence translation through the formation of secondary structures in mRNA. For this reason, we investigated whether incorporating information about structural interactions into the model could enhance its predictive power. To achieve this, we utilized predictions from the ArmNet model, which secured first place in the Ribonanza RNA 3D structure prediction challenge^72^. Predicted nucleotide-resolution reactivity profiles generated using the ArmNet model were incorporated into PARADE as additional input channels, providing sequence-specific structural information (see Extended Data Fig. 2D). During the prediction process, constant flanking regions from the plasmid (flanking 95 nts for both 5’ and 3’ ends) were added to each of the sequences.

### PARADE Diffusion model

To generate UTRs, we used a cold diffusion approach which we previously applied to designing yeast promoter sequences^37^. First, we trained the model to revert nucleotide substitutions in the sequences with known activity. To this end, random mutations were introduced into the sequences with the activity measured in MRPA Library 1. The number of mutations for each sequence was chosen from a uniform distribution, ranging from 0 to 200 for 5’ UTR and from 0 to 500 for 3’ UTR. The model received input consisting of six channels: four channels for one-hot-encoded nucleotides, one channel representing the number of mutations introduced into the sequence, and one channel for the activity of the original (non-mutated) sequence.

The cross entropy loss was calculated based on the similarity between the restored sequence and the reference sequence. Additionally, the Kullback–Leibler divergence loss was used to account for the GC content of the sequences (see Extended Data Fig. 4E). An independent model was trained for each cell line using the AdamW optimizer with a learning rate of 0.001, a batch size of 1024, and 1000 batches per epoch. Each dataset was split into training and validation sets at a ratio of 1:4.

Sequence generation then occurred through an iterative process of introducing sequence mutations to reach the target activity using the “cold diffusion” approach. The input to the model consists of a random one-hot encoded sequence, the number of mutations *n* (200 for the 5’ UTR and 500 for the 3’ UTR), and the desired expression level. Initially, a default number of mutations of *n*-1 is introduced into the starting sequence. The mutated sequence is then fed back to the model with *n*-1 specified in the ’mutation’ channel. This cycle continues until the number of ’uncorrected’ mutations reaches 0 where the activity of the sequence should have approached the target (n=50 and n=150 were used for 5’ and 3’ UTRs, respectively).

### Genetic Algorithm

As an additional approach to 5’- and 3’ UTR generation, we employed a genetic algorithm optimizing the predicted difference in activity between each particular pair of the cell lines. For implementation, we used PyGAD ^79^. The algorithm was launched for each ordered pair of the available cell lines with 10 different random seeds. The genetic algorithm was running for 25 generations of 10 000 sequences. In each cycle, the top half of sequences were selected to create ‘offsprings’ with adaptive mutation rates: 0.2 for sequences with the fitness below average and 0.05 for sequences with the fitness above average, and two-point crossovers with p=0.1. For the selection phase, we used a steady-state selection process, with top 50% of the sequences participating in forming the next generation, and one sequence with the highest fitness always saved in the population. As the fitness function, we used the difference between PARADE predictor outputs for the corresponding cell lines, with -10 added to 5’UTR sequences containing AUG effectively negating the model’s ability of creating uAUGs in the sequence pool.

### Selection of Sequences for Library 2

Library 2 was composed of sequences generated by several approaches: Diffusion models, Genetic Algorithm, motif seeding, random generation, and manually selected control sequences.

#### Dififiusion-Generated Sequences

An individual Diffusion model was trained for each cell line. For each cell line, we generated 1 million sequences, each targeting a specific expression value within the range of [2.0, 3.0]. For these sequences, the 4-mer composition score was calculated in the following manner. First, each 4-mer was scored by its occurrences in the native set against the dinucleotide shuffled set: 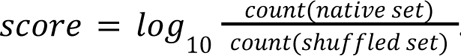. Second, the overall score for each sequence was defined as the sum of the scores for all 4-mers in the sequence. To filter out sequences with extreme scores, we removed 5’ UTR sequences with overall scores less than -3 or greater than 4.5, and 3’ UTR sequences with scores less than -5 or greater than 10. The remaining sequences were divided into five quintiles by the predicted average activity, and each group was further divided into five quintiles by the sequence entropy. Within each subgroup, sequences were ranked by the CTAD (Cell-Type Activity Difference) calculated sequentially for each pair of cell lines. For each pair, both negative and positive CTAD were considered to ensure the inclusion of sequences with higher activity in either cell line. From each subgroup, the top sequences were selected: 140 per group for 3′UTRs and 210 per group for 5′UTRs. Additionally, for each cell type, 300 sequences with uniformly distributed target activity between 2.0 and 3.0 were selected from the Diffusion-generated sequence pool.

#### Genetic Algorithm-Generated Sequences

Sequences were designed using a Genetic Algorithm to maximize CTAD for each pair of cell lines. Ten launches with different seeds were performed for each pair, resulting in 100,000 sequences per pair. These sequences were divided into quintiles by predicted average activity. Within each group, sequences were ranked by CTAD, and the top sequences were chosen for Library 2.

#### Motifi-Based Sequences

3000 sequences with the occurrences of identified RBP motifs were designed as described below under “Motif Analysis and Motif Group Sequence Constructionˮ.

#### Random Sequences

2,000,000 sequences were generated by randomly shuffling sequences from Library 1, preserving either mono- or di-nucleotide content (for 1 million each). Filtering was performed as described for Diffusion. Sequences were divided into quintiles by predicted average activity and further divided into quintiles by entropy. Within each subgroup, sequences were ranked by CTAD for each pair of cell lines, considering both directions, and the top sequences were selected: 70 per group for 3′UTRs and 98 per group for 5′UTRs.

#### Control Sequences

The control group included sequences from UTRs used in Pfizer’s COVID-19 vaccine BNT162b2^80^ and Moderna’s COVID-19 vaccine mRNA-1273^49^, 16 viral regulatory elements identified as driving mRNA stability^50^, and regulatory elements previously tested by Leppek^10^. Sequences longer than 50 nucleotides for 5’UTRs or 240 nucleotides for 3’UTRs were split into overlapping segments of the corresponding lengths. For 3’UTR sequences shorter than 240 nucleotides, padding was applied using a sequence that exhibited minimal activity changes across all cell lines in Library 1 (CCAUCCGCCAUUCCGACUGCUAAAAGCGAAUGUAGUCAGGCCCCUUUCAUGCUGUGAGACCUCCUGGAACACUGGCAUCUCUGAGCCUCCAGAAGGGGUUCUGGGCCUAGUUGUCCUCCCUCUGGAGCCCCGUCCUGUGGUCUGCCUCAGUUUCCCCUCCUAAUACAUAUGGCUGUUUUCCACCUCGAUAAUAUAACACGAGUUUGGGCCCGAAUCAGUGUGUUCUCAUCAUUUUUCAGG). This padding sequence was selected based on two criteria: (1) High coverage, defined as a median coverage of 500 reads across all libraries; and (2) Neutral activity, where activity was estimated as the weighted average of bin numbers (1–4) using normalized bin counts across replicates as weights. The activity value for this sequence deviates by no more than 0.05 from the “neutral activityˮ of 2.5 in every replicate for all cell lines.

### PARADE embeddings analysis

To quantitatively analyze the diversity of native, random, and generated sequences, we obtained their embeddings from the PARADE predictive model in the following manner. The model was dissected before the final convolutional layer, and the resulting tensor was averaged across each of the 32 channels. By supplying a sequence with different cell line markers in the input tensor to the network, the obtained cell line-specific embeddings were concatenated sequence-wise. In addition, k-mer frequencies for k=1, 2, or 3 were considered as extra features.

The embeddings were obtained for 1M random sequences generated by a 2-step model: nucleotide probabilities were taken from the Dirichlet distribution, and the sequences were generated as an i.i.d. series (Bernoulli model). The nucleotide probabilities sampling allowed for higher diversity of the generated sequences. Low-complexity sequences (measured by Shannon entropy of nucleotide frequencies) were filtered out of the distribution. The embeddings for the results of the genetic algorithm and diffusion model generation were extracted in the same way. An orthogonal basis in the latent space was selected by applying PCA to the standardized embedding of each of the randomly generated sequences and extracting principal coordinate axes, having other sequences projected onto them. Two of the components with the highest percentage of the explained variance were used for the visualization.

### Motif analysis and motif-seeded sequence construction

For motif analysis, we employed FIRE ^81^ and TF-MoDISco ^82^. **FIRE.** For FIRE, differentially active sequences between pairs of cell lines were identified using DESeq2^73^. For each pair of cell lines, the log fold change and adjusted p-value for each sequence were combined into a single value, vv, calculated as: *v* = *s* · (1 − *p*), where *p* is the adjusted P-value calculated by DESeq2, and *s* is the sign of the log-fold change between the two groups. Positive *v* values indicate activity favoring the first cell line, while negative *v* values indicate activity favoring the second cell line. The resulting *v* values were quantized into 50 bins and provided as input to FIRE for motif identification. To identify “activating” or “repressing” motifs, the activity of each sequence was quantified using the weighted average of the bin numbers (1-4), using the sum of normalized counts in the bin in all replicates as the bin weights. The center of mass values were then averaged across all cell lines, quantized into 50 bins, and provided to FIRE.

For constructing sequences containing multiple motifs associated with differential expression (referred to as “motif groups”), clusters of motifs were identified using FIRE. For each pair of cell lines, clusters of motifs that were preferentially active in one cell line over the other were selected. From each cluster, one representative motif was chosen randomly. Each sequence was designed to include a random combination of occurrences of these representative motifs, without replacement. For 5′ UTRs, sequences were designed to contain the occurrences of two motifs, while for 3′ UTRs, sequences were designed to contain the occurrences of four motifs. For each motif assigned to a sequence, a motif occurrence was randomly selected from those found in sequences in Library 1. The selected motif hit and its surrounding sequence were extracted, with a total length of 60 nucleotides for 3′UTRs and 25 nucleotides for 5′UTRs.

To construct sequences containing multiple activating or repressing motif occurrences, the same procedure was used, except that the clusters of motifs were obtained from FIRE analysis of generally activating or repressing motifs (defined by the center of mass described above) rather than motifs that differ between two cell lines.

#### TF-MoDISco

For TF-MoDISco, the input consisted of an array of genomic sequences (as TF-MoDISco uses the DNA alphabet) in one-hot encoding, an array of contribution scores where each position represented the importance of nucleotides corresponding to sequences in the previous array, and an array of hypothetical contribution scores obtained by *in silico* saturation mutagenesis.

Motifs were extracted from the TF-MoDISco output by filtering by information content with a threshold of 0.5. Motif finding was performed using SPRY-SARUS v.2.1.0^83^ with sum-occupancy scoring ^84^ with the parameters --naive --pfm-pseudocount 0.1. The resulting scores were employed for the partial Pearson correlation analysis between the presence of motif occurrences in the sequences (measured by the motif scanning score) and the activity levels of the sequences.

To reduce redundancy, the motifs were clustered by PWM similarity to each other and to known motifs from the oRNAment^85^ and CIS-BP-RNA (human subset) databases^86^. The similarity between motifs was quantified using the Jaccard index computed by the MACRO-APE^87^ EvalSimilarity function, employing the parameters --ppm -p 0.05 --position all,direct. In total, we clustered 927 and 2436 motifs from 5’UTR and 3’UTR respectively. Based on the Jaccard index, motifs were clustered using UPGMA (Unweighted Pair Group Method with Arithmetic Mean) clustering on the distance matrix with the cutoff of the number of clusters by the silhouette score of 0.16 and 0.098 and we obtained 350 and 1000 clusters for 5’UTR and 3’UTR respectively. For each cluster, a representative reference (known) motif was identified among the members of the cluster; otherwise, the most similar reference motif was selected with the similarity threshold of 0.25. If no known motifs with similarity higher than 0.25 were found, the motif was considered novel.

To investigate the impact of motifs on sequence activity controlling for the confounding effect of nucleotide composition, we employed the partial Pearson correlation (computed with Pingouin v0.5.4) between the motif scores and the measured activity. Partial correlation measures the degree of association between two variables while removing the influence of one or more additional variables, in our case, the nucleotide composition.

### T Cell and B Cell Culture

Peripheral blood mononuclear cells (PBMCs) were obtained from leukapheresis products (Leukopaks, STEMCELL Technologies). CD8+ T cells were isolated from the PBMCs using the EasySep™ Human CD8+ T Cell Isolation Kit (STEMCELL Technologies), which ensures high purity while preserving cell functionality. The isolated CD8+ T cells were resuspended in ImmunoCult™-XF T Cell Expansion Medium (STEMCELL Technologies), supplemented with Human Recombinant IL-2 (10 ng/mL, STEMCELL Technologies) to promote survival and proliferation. For activation, T cells were stimulated with ImmunoCult™ Human CD3/CD28 T Cell Activator (STEMCELL Technologies) at a final concentration of 25 µL/mL. Cultures were initiated at a density of 0.5–1 × 10⁶ cells/mL in a humidified incubator at 37°C with 5% CO₂. Cell density and viability were assessed every 2–3 days, and cultures were maintained by replenishing fresh medium supplemented with IL-2 to sustain optimal conditions. Expanded T cells were harvested after 7–10 days for downstream functional assays, including flow cytometry and cytotoxicity evaluation. B cells were isolated from the same leukapheresis products using the EasySep™ Human B Cell Isolation Kit (STEMCELL Technologies). Purified B cells were cultured in ImmunoCult™-XF B Cell Base Medium (STEMCELL Technologies), supplemented with ImmunoCult™-ACF Human B Cell Expansion Supplement to ensure consistent activation and expansion. B cells were seeded at a density of 0.5–1 × 10⁶ cells/mL and cultured in a humidified incubator at 37°C with 5% CO₂. Cultures were monitored every 2–3 days to assess cell density and viability. The medium was replenished with fresh complete medium to maintain nutrient and cytokine levels. Functional assays, including ELISA and flow cytometry, were performed post-expansion to evaluate B cell activity and phenotype.

### T Cell Electroporation

Expanded CD8+ T cells were subjected to RNA transfection using the 4D-Nucleofector® X Unit (Lonza) and the Human T Cell Nucleofector® Kit (Lonza). Prior to electroporation, cells were harvested, washed twice with phosphate-buffered saline (PBS), and resuspended in the electroporation solution provided in the kit at a final density of 1 × 10⁷ cells/mL. Then, the mRNA encoding the target protein was added to the cell suspension at a final concentration of 10–15 µg per 10⁶ cells, ensuring optimal transfection efficiency. The mixture was transferred into a nucleocuvette™ provided with the kit. Electroporation was performed using the pre-optimized program EO-115, specifically tailored for human T cells, on the 4D-Nucleofector® X Unit. Immediately after electroporation, cells were gently transferred to pre-warmed ImmunoCult™-XF T Cell Expansion Medium supplemented with Human Recombinant IL-2 (10 ng/mL) and incubated in a humidified incubator at 37°C with 5% CO₂. Cell viability and recovery were assessed 24 hours post-electroporation using a hemocytometer and trypan blue exclusion assay. Electroporated T cells were cultured for 24–72 hours to allow for RNA translation and protein expression.

### Combinatorial treatment of cells with BSO and mRNA

HepG2 cells were cultured in Dulbecco’s Modified Eagle Medium (DMEM; Gibco) supplemented with 10% fetal bovine serum (FBS; Gibco), 1% penicillin-streptomycin (Gibco), and 2 mM L-glutamine. Cells were maintained at 37°C in a humidified incubator with 5% CO₂. For experiments, cells were seeded in 96-well plates at a density of 5 × 10³ cells per well and allowed to adhere overnight. Primary human T cells were isolated and cultured as described before. For HepG2 cells, the glutathione-depleting agent L-Buthionine-sulfoximine (BSO; Sigma-Aldrich) was prepared in sterile PBS and added to cells at a final concentration of 0.5–5 mM, depending on the experimental condition. RNA transfection into HepG2 cells was performed using the TransIT®-mRNA Transfection Kit (Mirus Bio) according to the manufacturer’s instructions. Briefly, RNA was diluted in Opti-MEM™ (Gibco), mixed with the TransIT-mRNA reagent, incubated for 3–5 minutes at room temperature, and added to cells at a final RNA concentration of 1–2 µg/mL. For T cells, BSO was added at similar concentrations, and RNA was introduced via electroporation as described. Cell viability was assessed using the CellTiter-Glo® Luminescent Cell Viability Assay (Promega). At the end of the treatment period (24–72 hours), plates were equilibrated to room temperature. An equal volume of CellTiter-Glo® reagent was added to each well, and the plates were mixed on an orbital shaker for 2 minutes to ensure cell lysis. Luminescence was measured using a plate reader (BioTek Synergy HTX), with results expressed as relative luminescence units (RLUs).

### Pooled RNA stability measurements in T cells

49 3’ UTRs were randomly selected from 1000 3’ UTRs with the highest PARADE-predicted effects on mRNA stability in Jurkat cells. To assess the stability of pooled RNAs, multiple mRNAs were mixed in equal ratios with lipids and encapsulated into LNPs. After 48 hours of incubation, LNPs were removed by replacing the culture medium with fresh, LNP-free medium. Samples were collected for RNA extraction at defined time points (post-LNP removal). For the RNA extraction, cells were harvested, and RNA was extracted using the TRIzol™ reagent (Thermo Fisher Scientific) and ethanol precipitation protocol. Briefly, 1 mL of TRIzol™ reagent was added to each cell sample, and phase separation was performed by adding chloroform and centrifuging. The aqueous phase was collected, and RNA was precipitated with ethanol, washed, and resuspended in nuclease-free water. cDNA synthesis was performed using Thermo Scientific Maxima Reverse Transcriptase (primer CTCTTTCCCTACACGACGCTCTTCCGATCTNNNNNNNNNNNggcagaatccagatgctcaagg), and the UTRs were amplified by targeted PCR with NEB Q5 polymerase (primers TCTCAGTACAGCCACCAAGGAC and CTCTTTCCCTACACGACGCTCTTC). The amplified libraries were purified using Zymo Research MagBinding Beads and sequenced on a NovaSeq X platform at the UCSF Center for Advanced Technology. Sequencing reads were processed by trimming adapter sequences with cutadapt^75^, extracting Unique Molecular Identifiers (UMIs) with umi-tools^76^, and aligning the reads to the reference set of sequences using bwa mem^77^. RNA stability was estimated as the ratio of read counts in samples collected on day 3 to the read counts in the input RNA library.

### Luciferase Measurement in HepG2 Cells, T Cells, and B Cells

NanoLuc and Firefly luciferase activities were quantified using the Nano-Glo® Dual-Luciferase® Reporter Assay System (Promega) following the manufacturer’s protocol. HepG2 cells, T cells, and B cells were cultured under appropriate conditions and treated according to the experimental design. After treatment, cells were lysed using Passive Lysis Buffer provided in the assay kit. Lysates were transferred to a white, opaque 96-well microplate to enhance luminescence signal detection. Firefly luciferase activity was measured by adding 50 µL of Luciferase Assay Reagent II (LAR II) to each well containing 50 µL of lysate. The plate was incubated at room temperature for 2 minutes to allow the reaction to stabilize, and luminescence was recorded using a microplate luminometer. To measure NanoLuc luciferase activity from the same wells, 50 µL of Stop & Glo® Reagent (containing Nano-Glo® Substrate) was added directly to each well. This reagent quenched the Firefly luciferase signal while simultaneously activating NanoLuc luciferase. After a 3-minute incubation at room temperature, NanoLuc luminescence was measured using the same luminometer. Relative luminescence units (RLUs) for both Firefly and NanoLuc luciferase activities were recorded and normalized to background luminescence, which was measured from wells containing only reagents without lysates. The total protein output was calculated as the Area Under the Curve (AUC) of luciferase measurements across time points.

### mRNA Synthesis and Purification

The UTRs of interest, along with coding region of interest (Luciferase, CD19 CAR, P16, or PTEN) was cloned into a pEZ vector (Epoch Life Science), containing the T7 RNA Polymerase promoter and a segmented poly(A) tail ^88^. Linearized DNA templates encoding the gene of interest were prepared and purified to ensure efficient transcription. Following linearization, DNA was cleaned using the Zymo Research DNA Clean & Concentrator-25 Kit (Zymo research) according to the manufacturer’s protocol. Then, mRNA was synthesized using the Takara IVTpro T7 mRNA Synthesis Ki (Takara Bio) following the manufacturer’s protocol. DNA templates were used in a transcription reaction with a total volume of 20–50 µL, including linearized and purified DNA, NTP mix (provided in the kit), and the Takara transcription enzyme mix. For mRNA capping, the Clean Cap Reagent (TriLink BioTechnologies, Cat #N-7113) were incorporated during the transcription reaction to produce 5’ capped mRNA. The CleanCap system was added to the IVT reaction in the proportions recommended by Takara, ensuring high-efficiency co-transcriptional capping. The reaction was incubated at 37°C for 2 hours to facilitate complete mRNA synthesis and capping. Following the IVT reaction, the synthesized mRNA was purified using the Zymo Research RNA Clean & Concentrator-25 Kit (Zymo research) to remove residual enzymes, unincorporated nucleotides, and contaminants. The purification procedure was carried out as per the manufacturer’s instructions. The purified mRNA was analyzed for Integrity and size were verified using the Agilent TapeStation system (Agilent Technologies). Results from the TapeStation provided clear confirmation of RNA integrity and appropriate fragment size distribution.

### LNP preparation for *In Vivo* and T cells

The lipid mixture for *In vivo* was prepared using DLin-MC3-DMA (Cayman Chemical), 1,2-DSPC (Cayman Chemical), cholesterol (Cayman Chemical), and DMG-PEG (2000) (Cayman Chemical) with the following molar ratios: 50:10:38.5:1.5, respectively. Each component was dissolved in absolute ethanol to prepare the stock solutions. The prepared lipid mixture was combined with RNA solutions to generate LNPs using the NanoAssemblr™ Ignite™ nanoparticle formulation systems (Precision NanoSystems) Using a flow rate of 9 mL/min and an aqueous-to-organic phase flow rate ratio of 2:1, the mixture was subsequently dialyzed overnight against TBS buffer (20 mM Tris, 0.9% NaCl, pH 7.4). The RNA solution (The mRNA was diluted in 100 mM sodium acetate buffer (pH 4)) and lipid mixture were loaded into the NanoAssemblr Ignite using Ignite+™ cartridges, which utilizes microfluidic mixing to ensure precise and reproducible nanoparticle formation. The system parameters, such as flow rate and total volume, were adjusted as per the Ignite protocol to optimize particle size and encapsulation efficiency. The freshly prepared LNPs were purified to remove unencapsulated RNA, ethanol, and other impurities using Amicon Ultra-15 centrifugal filters (MilliporeSigma) with a molecular weight cutoff (MWCO) of 10 kDa. For purification, LNPs were diluted with nuclease-free PBS (pH 7.4) to a working volume of 15 mL and transferred into Amicon Ultra-15 filters. Then, the samples were centrifuged at 4,000 × g for 10–15 minutes at 4°C, reducing the volume to approximately 1 mL. The remaining LNPs were resuspended in PBS and the process was repeated 2–3 times to ensure thorough removal of ethanol and other contaminants. After purification, the LNPs were collected, and their concentration was measured using the RiboGreen RNA quantitation assay (ThermoFisher Scientific, Cat# R11490) following the manufacturer’s instructions. LNP size and polydispersity index were measured using dynamic light scattering (DLS), and encapsulation efficiency was assessed through RNA quantification via the RiboGreen assay. For the T cells, LNPs formulated using the GenVoy-ILM™ T cell kits for mRNA (Cytiva). LNP preparation was performed using the NanoAssemblr. Then, the mRNA encoding the target protein (1 mg/mL stock) was encapsulated in LNPs using the GenVoy-ILM T Cell Lipid Mix (Cytiva). Formulations were prepared by combining the mRNA aqueous phase with the lipid phase in the NanoAssemblr® Ignite Instrument at a flow rate ratio of 2:1 following the manufacturer’s protocol. The prepared LNPs were filtered, characterized for size and encapsulation efficiency, and diluted in sterile PBS, as described before. For transfection, 0.5 × 10⁶ T cells in 1 mL of complete medium were treated with 1 µg/mL apolipoprotein (Cytiva) and 3 µg of mRNA-loaded LNPs per 10⁶ cells. The cells were incubated with the LNPs at 37°C in a humidified incubator with 5% CO₂ for 48 hours to allow for mRNA uptake, translation, and protein expression.

### In Vivo patient-derived xenograft tumor implantations

Tumor fragments from a glioblastoma (GBM) patient-derived xenograft (PDX) model (GBM6) null for p16 were obtained from the brain tumor center preclinical therapeutic testing core at UCSF for implantation. Approximately 50 µL of homogenized tumor tissue was implanted into both flanks of 6–8-week-old nude (NU/J) mice (strain no. 002019, The Jackson Laboratory) using 18-gauge syringes. Tumors were allowed to establish over one week before treatment began. Mice were randomly assigned into four experimental groups to ensure unbiased treatment allocation. LNPs encapsulating p16 mRNA were prepared at a dose of 2 mg/kg mRNA. The LNPs were administered via intratumoral injection into each tumor site for five treatment rounds administered on days 1, 3, 5, 7, and 9, starting one week after the tumor injection. Tumor volumes in mice were measured using digital calipers to assess tumor growth and response to treatment. Mice were gently restrained, and if required, anesthetized with isoflurane to minimize stress and ensure accurate measurements. Tumor dimensions, including the width and length, were recorded in millimeters. The width was defined as the shortest diameter of the tumor, and the length was defined as the longest axis perpendicular to the width. Tumor volume was calculated using the formula: width squared multiplied by the length, divided by two.

### Intracerebral Tumor Establishment in Athymic Mice

Five- to six-week-old female athymic nu/nu homozygous mice (Envigo Laboratories, Livermore, CA) were housed under aseptic conditions and received intracranial tumor cell injections as previously described ^89^ and approved by the University of California San Francisco Institutional Animal Care and Use Committee. Briefly, mice were anesthetized via a combination of intraperitoneal injection of a mixture containing ketamine (100 mg/kg) and xylazine (10 mg/kg), along with inhalation of isoflurane. Then, 3 μL of a tumor cell suspension (300,000 cells) was injected into the right caudate putamen using a freehand method.

### Procedure for Intracerebral Cell Injection

All procedures were carried out under sterile conditions. Mice were anesthetized via a combination of intraperitoneal injection of a ketamine and xylazine mixture and inhalation of isoflurane. The scalp was surgically prepped, and a ∼10 mm incision was made over the frontal to parietal bone. The skull surface was exposed, and a small hole was created 3.0 mm to the right of the bregma and just anterior to the coronal suture using a 25-gauge needle. A 26-gauge needle attached to a Hamilton syringe was inserted into the hole. The needle was fitted with a sleeve to limit the injection depth to 3-4 mm. A 3 μL cell suspension was injected slowly (∼1 μL/min) using the freehand method. After injection, the needle was removed, and the skull surface was swabbed with hydrogen peroxide before sealing the hole with bone wax to prevent reflux. The scalp was closed with surgical staples.

### Bioluminescence Monitoring of Intracranial Tumor Growth

For bioluminescence imaging (BLI), mice were anesthetized using inhalation of isoflurane and administered 150 mg/kg of luciferin (D-luciferin potassium salt, Gold Biotechnology, St. Louis, MO) via intraperitoneal injection. Ten minutes after luciferin administration, mice were examined for tumor bioluminescence using an IVIS Lumina imaging station and Living Image software (Caliper Life Sciences, Alameda, CA). Regions of interest were quantified as photons per second per steradian per square centimeter ^89^.

### Convection-Enhanced Delivery

Our approach was similar to that previously described ^90^. Briefly, infusion cannulae were fabricated using silica tubing (Polymicro Technologies, Phoenix, AZ) fused to a 0.1 mL syringe (Plastic One, Roanoke, VA) with a 0.5 mm stepped-tip needle protruding from the silica guide base. The syringe was loaded with RNA-LNPs and attached to a microinfusion pump (Bioanalytical Systems, Lafayette, Ind.). The syringe and silica cannula were lowered through a puncture hole in the skull ^89,91^ into the same region of the caudate putamen where tumor cells were previously injected. RNA-LNPs was infused at a rate of 1 μL/min until a total volume of 15 μL was delivered. Cannulae were removed 2 minutes after the infusion was completed. The LNPs were administered over a period of three weeks, with a single dose given each week via CED.

### In Vivo Luciferase mRNA Delivery

Female NOD-scid gamma (NSG) mice (strain no. 005557, The Jackson Laboratory), aged 4–6 weeks, were used for the in vivo mRNA delivery experiments. All animal procedures were approved by the institution’s Animal Care and Use Committee (IACUC). F-luciferase lipid nanoparticles (F-luciferase LNPs) containing 0.5 mg/kg luciferase mRNA were prepared for injection. Mice were administered the LNPs via intravenous injection into the tail vein. For the tissue-specificity experiment, 24 hours post-injection, mice received an intraperitoneal injection of D-luciferin (Promega) at a dose of 150 mg/kg. Following D-luciferin administration, mice were sacrificed, and their livers and spleens were harvested for imaging. Bioluminescence detection was performed using the IVIS Spectrum Imaging System (PerkinElmer). For the RNA stability experiment, mice received an intraperitoneal injection of D-luciferin (Promega) at a dose of 150 mg/kg. Following D-luciferin administration, mice were anesthetized in a chamber using 3% isoflurane (Piramal Healthcare Limited) and positioned on the imaging platform while anesthesia was maintained at 2% isoflurane via a nose cone. Imaging was performed 5 minutes after D-luciferin administration. Images were analyzed using Living Image Software (PerkinElmer). The total radiant efficiency (photons/sec/cm²/sr) for each organ was quantified and normalized to the total area of the organ imaged.

### Harvesting p16 Tumors and RNA Quantification via RT-qPCR

Tumor samples were harvested and homogenized using a tissue homogenizer to ensure complete disruption of the tissue. Total RNA was extracted using TRIzol reagent (Thermo Fisher Scientific) following the manufacturer’s protocol, incorporating ethanol precipitation for RNA purification. The extracted RNA was quantified, and cDNA was synthesized using the Maxima H Minus Reverse Transcriptase kit (Thermo Fisher Scientific) with random primers (Thermo Fisher Scientific) according to the recommended procedure. Quantitative PCR (qPCR) was performed using SYBR Green PCR Master Mix (Thermo Fisher Scientific), with human *CDKN2A* (Cyclin-Dependent Kinase Inhibitor 2A) as the target gene and human *HPRT1* (Hypoxanthine Phosphoribosyltransferase 1) as the housekeeping gene. Gene expression levels were normalized to HPRT and analyzed using the ΔΔCt method.

## Authors Contributions

M.K., I.K., and H.G. designed the study. M.K. developed the reporters for massively parallel assays. M.K., S.L.,T.M., and K.M. performed massively parallel reporter assays. A.Z. and E.A. under the supervision of D.P. and I.K. developed PARADE Predictor and PARADE Generator. A.Z., E.A., and M.K. performed data analysis and designed the sequence libraries. M.K., H.Y. and K.M. performed RNA activity measurements in cells. H.Y. and D.R.R. performed RNA activity measurements in animals. M.K., I.K. and H.G. wrote the manuscript with input from all authors.

## Data and Code Availability

Sequencing data has been deposited in the Gene Expression Omnibus. The implementation of the PARADE framework is available at GitHub: https://github.com/autosome-ru/parade.

## Acknowledgments

We thank Koh, Kyung Duk for help with the 5’UTR reporter design. We thank Brian Plosky and Chiara Ricci-Tam for thoughtful comments. We thank Ilya Vorontsov for his assistance with RNA motifs clustering.

This study was supported in part by HDFCCC Laboratory for Cell Analysis Shared Resource Facility through a grant from NIH (P30CA082103). A.Z. was supported by a personal fellowship from the Non-commercial Foundation for Support of Science and Education ‘INTELLECT’.

## Competing Interests

M.K. and H.G. are inventors on a provisional patent related to this study. The remaining authors declare no competing interests.

## Notes

### Competing Interest Statement

The authors have declared no competing interest.

